# Aurora B phosphorylates Bub1 to promote spindle assembly checkpoint signaling

**DOI:** 10.1101/2021.01.05.425459

**Authors:** Babhrubahan Roy, Simon J. Y. Han, Adrienne N. Fontan, Soubhagyalaxmi Jema, Ajit P. Joglekar

## Abstract

Accurate chromosome segregation during cell division requires amphitelic chromosome attachment to the spindle apparatus. It is ensured by the combined activity of the Spindle Assembly Checkpoint^1^ (SAC), a signaling mechanism that delays anaphase onset in response to unattached chromosomes, and an error correction mechanism that eliminates syntelic attachments^2^. The SAC becomes active when Mps1 kinase sequentially phosphorylates the kinetochore protein Spc105/KNL1 and the signaling proteins that Spc105/KNL1 recruits to facilitate the production of the Mitotic Checkpoint Complex^3-8^ (MCC). The error correction mechanism is regulated by the Aurora B kinase, but Aurora B also promotes SAC signaling via indirect mechanisms^9-12^. Here we present evidence that Aurora B kinase activity directly promotes MCC production by working downstream of Mps1 in budding yeast and human cells. Using the ectopic SAC activation (eSAC) system, we find that the conditional dimerization of Aurora B in budding yeast, and an Aurora B recruitment domain in HeLa cells, with either Bub1 or Mad1, but not the phosphodomain of Spc105/KNL1, leads to ectopic MCC production and mitotic arrest^13-16^. Importantly, Bub1 must recruit both Mad1 and Cdc20 for this ectopic signaling activity. These and other data show that Aurora B cooperates with Bub1 to promote MCC production, but only after Mps1 licenses Bub1 recruitment to the kinetochore. This direct involvement of Aurora B in SAC signaling may maintain SAC signaling even after Mps1 activity in the kinetochore is lowered.

## Results & Discussion

### Dissecting the contributions of Bub1 and Mad1 in Mps1-driven Mitotic Checkpoint Complex (MCC) formation

To activate the SAC and delay anaphase onset, unattached kinetochores produce the MCC, which is a complex of four proteins: the ‘closed’ form of Mad2, Cdc20, BubR1, and Bub3. MCC production is licensed by the Mps1 kinase, which sequentially phosphorylates SAC proteins within the kinetochore to promote their recruitment^3-6^ (Figure 1A). Cdc20 is recruited by Bub1 and BubR1 through constitutive interactions^17^. SAC proteins recruited interact to produce the MCC. Although Mps1 plays the dominant and essential role in this process, Aurora B kinase activity is also required for maximal signaling^9-11,18-23^. In budding yeast, Aurora B phosphorylates Mad3/BubR1^24^. Whether it phosphorylates other SAC signaling proteins to catalyze MCC production has been difficult to determine, mainly because Mps1 and Aurora B act concurrently within unattached kinetochores. Therefore, to study the effect of Aurora B on SAC proteins, we used the ectopic SAC activation assay or “eSAC”^13^. In this assay, a mitotic kinase domain is conditionally dimerized with a cytosolic phosphodomain of the kinetochore protein Spc105/KNL1 or Bub1^13-16^. The ensuing signaling activity is evident as a mitotic delay.

**Figure 1.**
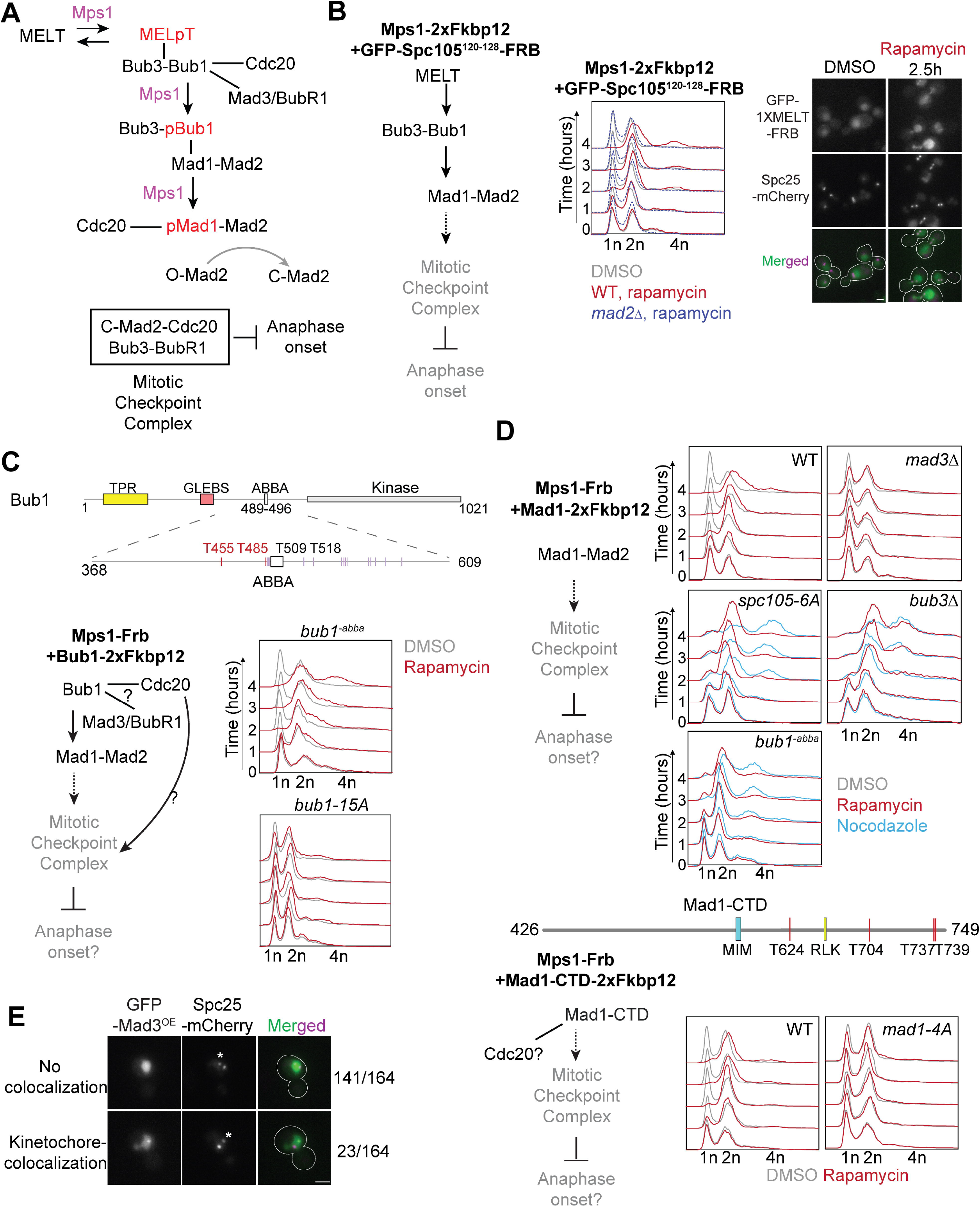
Dissection of the regulatory role of the Mps1 kinase in the SAC signaling cascade using the ‘eSAC’ system. (A) A simplified schematic of the SAC signaling cascade in budding yeast. Black arrows represent regulated recruitment of downstream signaling proteins; black lines represent constitutive protein-protein interactions. The ‘Mps1’ label over an arrow signifies phosphoregulation of the protein recruitment step by Mps1. Gray arrow represents the assembly of sub-complexes into the mitotic checkpoint complex. (B) The ectopic activation of the SAC signaling cascade (simplified on the left hand) by the rapamycin-induced dimerization of Mps1-2xFkpb12 with a cytosolic fragment of Spc105 containing just one MELT motif (GFP-Spc105^120-128^-Frb). Graphs on the right show the quantitation of cellular DNA content using flow cytometry over 4 hours after the addition of rapamycin to the growth media. Strain genotype is indicated at the top of each graph. Grey, red and dashed blue lines show cytometry results for DMSO-treated, rapamycin-treated wild-type and *mad1Δ* cells respectively. Representative micrographs of yeast cells expressing the indicated, fluorescently labeled proteins (right). Notice that the cytosolic GFP-Spc105^2-128^-Frb displays faint kinetochore colocalization in rapamycin-treated cells, likely because Mps1 localizes to bioriented kinetochores^14^. Scale bar∼3.2µm. (C) Top: Domain organization of full length Bub1 (Bub1^FL^) and its middle domain in budding yeast. Bottom left: Schematic of the potential effects of the rapamycin-induced dimerization of Mps1-Frb with Bub1-2xFkbp12. Bottom right: Graphs show flow cytometry of DMSO-treated (black lines) and rapamycin-treated (red lines) cells with the indicated genotype. (D) Top: Flow cytometry panels showing effects of rapamycin-induced dimerization of Mps1-Frb with the Mad1-2xFkbp12 in wild-type, in absence of Mad3, in presence of spc105-6A, in absence of Bub3 and in presence of bub1^-abba^. Plots with black, red and cyan lines indicate cells treated with DMSO (control), rapamycin and nocodazole respectively. Middle: Domain organization of Mad1-CTD (amino acid 426-749) in budding yeast. Bottom: In left, a partial schematic of SAC cascade is shown. In right the potential effects of the rapamycin-induced dimerization of Mps1-Frb with the Mad1-CTD-2xFkbp12 or Mad1-CTD^4A (T624A, T704A, T737A, T739A)^-2xFkbp12 are shown. Color scheme as in B-C. (E) Localization of ectopically expressed GFP-Mad3 in cells arrested in mitosis due to nocodazole treatment. Note that in nocodazole-treated yeast cells typically contain two kinetochore clusters. The larger cluster is proximal to the spindle pole bodies (not visualized), and kinetochores within this cluster are attached to short microtubule stubs. The smaller cluster is distal to the spindle pole bodies (asterisk), and the kinetochores within this cluster are unattached^58^. Scale bars ∼ 3.2 µm. See also Figure S1, Table S1, and Data S1A.

We first used the eSAC assay to clearly delineate the known Mps1 roles in SAC signaling relying on flow cytometry to quantify the DNA content of yeast cells (Figure 1B). As positive controls, we dimerized Mps1-2xFkbp12 with either ‘MELT’-motif containing fragments of the Spc105 phosphodomain or Bub1 or Bub3 fused to Frb and GFP by adding rapamycin to the growth media (Figure 1B). In all cases, cell populations shifted to 2n ploidy indicating a G2/M arrest^25^ (Figure 1B-C, Figure S1A-B, rapamycin panels in red). Cells lacking Mad2 did not arrest following rapamycin treatment (Figure 1B, dashed blue panel). Thus, the arrest required a functional SAC.

Bub1 contributes two activities for MCC formation. It recruits Mad1 via its central domain, and in human cells, Cdc20 via a conserved ‘ABBA’ motif^3,5,17,26-28^ (Figure 1C). To separate their contributions, we generated bub1^-abba^ wherein the hydrophobic residues predicted to interface with Cdc20 are mutated^26^. Induced dimerization of Mps1 with bub1^-abba^ caused a mitotic arrest suggesting that the ABBA motif is dispensable in Mps1-driven signaling (Figure 1C). Consistently, *bub1*^*-abba*^ cells arrested in mitosis following nocodazole treatment (Figure S1C). The fraction of cells escaping the mitotic block was higher compared to wild-type cells indicating that the SAC is weaker (Figure S1C). Mad1-mCherry recruitment to unattached kinetochores in *bub1*^*-abba*^ cells was also ∼50% lower^28^ (Figure S1D). Bub1-Cdc20 interaction has been difficult to confirm using biochemical methods in HeLa cells^29^. We also could not confirm this interaction in yeast using immunoprecipitation, yeast two-hybrid assays, or microscopy (data not shown). However, the conservation of the ABBA motif sequence supports our assumption that bub1^-abba^ cannot bind Cdc20.

Mps1 phosphorylates many sites within the Bub1 central domain to promote the Bub1-Mad1 interaction^3,5,6,28,30^. To find the essential phosphorylatable residues, we generated numerous Bub1 mutants wherein subsets of the phosphorylation sites are non-phosphorylatable (Table S1). Some of these mutants weakened, but did not abolish, SAC signaling (Table S1). Mad1 localization to unattached kinetochores was also diminished, but not abolished, even in those mutants that significantly weakened the SAC (Figure S1D). Consistently, Mps1 dimerization with these phosphomutants led to robust eSAC signaling (Table S1). Finally, we dimerized Mps1 with bub1-15A, wherein 15 phosphorylation sites are non-phosphorylatable^3,16^. bub1-15A localized to unattached kinetochores in nocodazole-treated cells better than wild-type Bub1. However, it did not activate the either SAC or the eSAC (Figure 1C, Figure S1C and S1E, flow cytometry right panel). Thus, eSAC signaling driven by Mps1-Bub1 dimerization requires Bub1-mediated Mad1 recruitment.

In human cells, Mps1 phosphorylates Mad1 to facilitate the formation of Cdc20:C-Mad2^5,6^. Consistently, induced dimerization of Mps1 with Mad1 arrested yeast cells in mitosis. This arrest was not observed in *mad3Δ* cells, indicating that it was mediated by ectopic SAC signaling (Figure 1D). eSAC activity persisted in *spc105-6A* cells, wherein all six MELT motifs in Spc105 are non-phosphorylatable, and in *bub3Δ* cells. Thus, this ectopic SAC activation does not require Bub1 localization to the kinetochore. The latter observation also indicates that the budding yeast MCC can be formed ectopically without Bub3^31^. Consistent with the dispensability of the Bub1 ABBA motif for Mps1-driven eSAC signaling, Mps1-Mad1 dimerization produced a robust mitotic arrest in *bub1*^*-abba*^ cells (Figure 1D). Finally, Mps1 dimerization with mad1-4A, wherein residues implicated in the Mad1-Cdc20 interaction are non-phosphorylatable (Figure 1D), did not affect the cell cycle^5,6^. Interestingly, Mps1-Mad1 dimerization did not affect cell cycle progression in cells expressing *bub1-15A* likely because Bub1 scaffolding of Mad1 and Cdc20 is necessary for the ectopic MCC assembly^32,33^ (Figure S1F).

MCC formation also requires BubR1, known as Mad3 in budding yeast. In metazoa and fission yeast, BubR1 is recruited to the kinetochore by Bub1^16,34,35^. However, this Bub1-BubR1 interaction is likely not conserved in budding yeast^36^. To confirm this, we fused GFP to the N-terminus of Mad3 (C-terminal fusion of GFP to Mad3 makes it incompetent in SAC signaling, data not shown) and visualized it in nocodazole-treated cells. GFP-Mad3 expressed from the endogenous promoter was undetectable at unattached kinetochores (data not shown). Even when exogenous GFP-Mad3 was significantly overexpressed, it rarely colocalized with unattached kinetochores (Figure 1E and Figure S1G).

These observations suggest the following model for Mps1-driven SAC signaling in budding yeast. As established by previous studies, Mps1 licenses the sequential recruitment of Bub1-Bub3 and Mad1-Mad2 by phosphorylating the MELT motifs and Bub1. ABBA motif in Bub1 contributes to SAC signaling, but it is not essential. Mps1 also phosphorylates Mad1 to promote its interaction with Cdc20, and this is essential for MCC formation. Finally, the lack of GFP-Mad3 localization suggests that unattached yeast kinetochores mainly catalyze Cdc20-C-Mad2 formation.

### Testing whether Aurora B/Ipl1 kinase activity promotes MCC formation

The Aurora B kinase, known as Ipl1 in budding yeast, is implicated in SAC signaling. Ipl1 is positioned in unattached kinetochores to act on SAC proteins recruited to the kinetochore by Mps1 activity^37^ (Figure 2A). Therefore, it may also phosphorylate Spc105/KNL1, Bub1, or Mad1 and thus promote MCC formation. To test this hypothesis, we conditionally dimerized Ipl1 with either a fragment of Spc105 spanning its six ‘MELT’ motifs, Bub1, or Mad1. Rapamycin-induced dimerization of Ipl1 with the Spc105 fragment did not affect the cell cycle, but its dimerization with either Bub1 or Mad1 induced a G2/M arrest (Figure 2B). The morphology of these cells was consistent with a metaphase-arrest: large buds and an intact spindle with two bioriented kinetochore clusters (Figure 2B). Ipl1-Bub1 dimerization did not affect cell cycle progression in *mad1Δ* mutants (Figure 2B blue curve in the middle panel). Thus, the arrest required a functional SAC.

**Figure 2.**
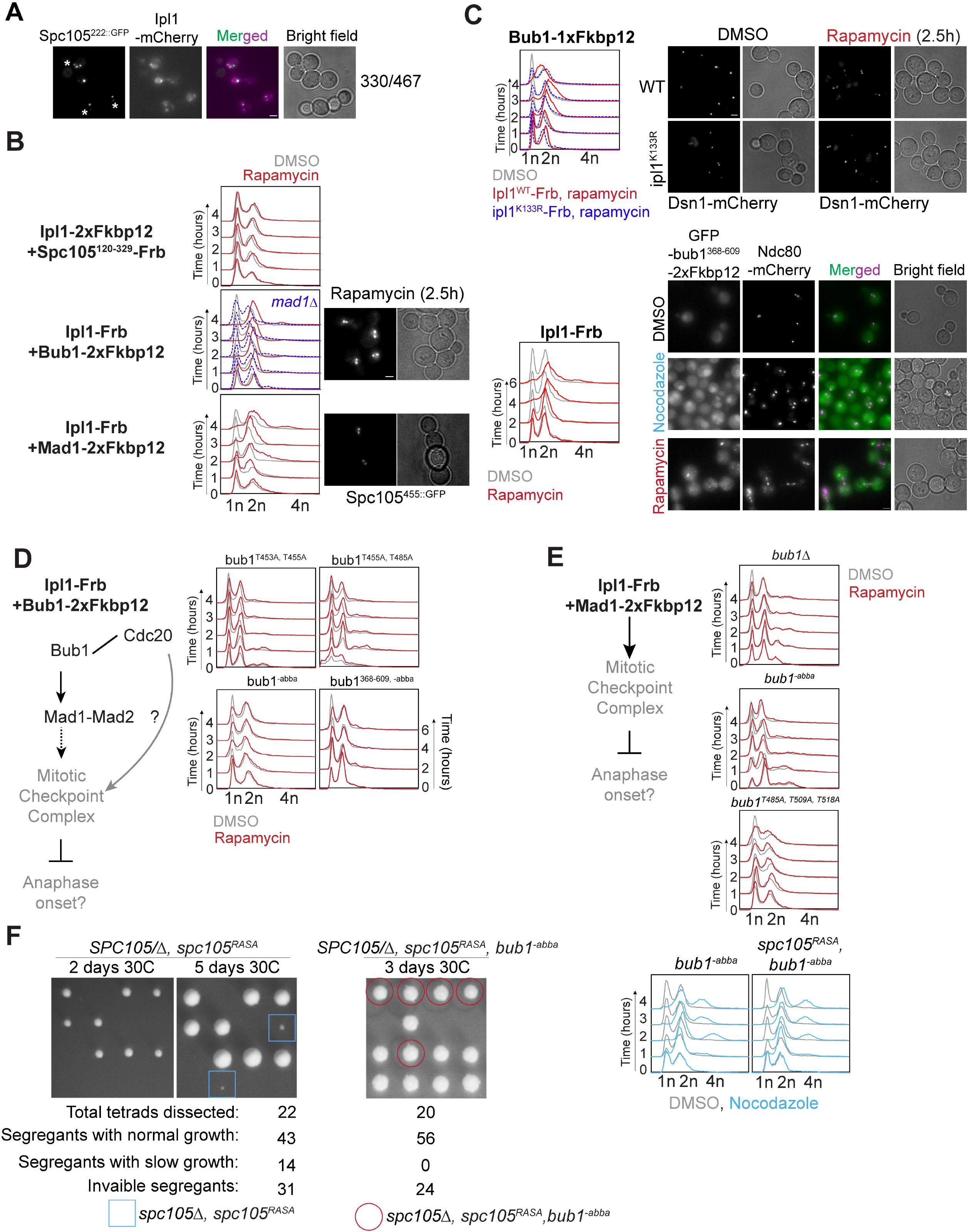
Rapamycin-induced dimerization of Aurora B/Ipl1 with Bub1 and Mad1, but not MELT motifs, leads to ectopic SAC activation. (A) Localization of Ipl1-mCherry and yeast kinetochores (visualized with Spc105^222::GFP^) in yeast cells arrested in mitosis due to nocodazole treatment. Asterisks mark the cluster of unattached kinetochores in each cell. Scale bar ∼ 3.2µm. (B) Left: Potential effects of the rapamycin-induced dimerization of Ipl1-Frb with the indicated SAC signaling protein. Middle: flow cytometry analysis of DMSO or Rapamycin-treated cultures of indicated strains. Color scheme as in Fig. 1. Right: Micrographs showing morphology of rapamycin-treated cells expressing Ip1-Frb and either Bub1-2XFkbp12 or Mad1-2XFkbp12. Scale bar∼3.2µm. (C) Top left: Flow cytometry analysis following rapamycin induced dimerization of Bub1-1xFkbp12 with Ipl1^WT^ (red) or ipl1^K133R^-Frb-GFP (dashed blue). Top right: Representative micrographs of yeast cells expressing Dsn1-mCherry after treatment with DMSO or rapamycin. Scale bar∼3.2µm. Bottom left: Flow cytometry panel showing effects of rapamycin induced dimerization of Ipl1-Frb with GFP-bub1^368-609^-2xFkbp12 (Color scheme: DMSO, grey; WT, red; -abba, dashed blue). Bottom right: Representative micrographs of yeast cells expressing Ndc80-mCherry and GFP-bub1^368-609^-2xFkbp12 after treatment with DMSO, rapamycin and nocodazole. Scale bar∼3.2µm. (D) Dissection of the contributions of Bub1-mediated recruitment of Mad1 and Cdc20 in ectopic SAC signaling driven by Mps1 using Bub1 point mutants. 2xFkbp12-tagged Bub1 mutants (indicated at the top of each flow cytometry panel) expressed from the genomic Bub1 locus. (E) Flow cytometric analysis of the effect of rapamycin induced dimerization of Ipl1 and Mad1 in absence of Bub1 (*bub1Δ*, top), in presence of bub1^-abba^ (middle) and in presence of bub1^T485A, T509A, T518A^ (bottom) on the cell cycle. (F) Left: Tetrad dissection of diploids with the indicated genotype. The plate with *SPC105/Δ, spc105*^*RASA*^ were imaged after 2 days (left) and 5 days (middle) before replica plating. The plate with *SPC105/Δ, spc105*^*RASA*^, *bub1*^*-abba*^ was replica plated after 2 days of incubation and then imaged on the 3^rd^ day. The growth rate of segregants expressing *spc105Δ, spc105*^*RASA*^, *bub1*^*-abba*^ (red circles) is similar to wildtype segregants. The numbers of inviable segregants for *spc105Δ, spc105*^*RASA*^, *bub1*^*-abba*^ and inviable or micro-colony forming segregants (blue squares) for *spc105Δ, spc105*^*RASA*^ are consistent with the probabilities predicted by the random segregation of three and two genes (0.33 and 0.5) respectively. Right: Flow cytometric analysis of cell cycle progression in nocodazole-treated cells carrying the indicated mutations. See also Figure S2, Table S1, Table S2, Data S1B.

To ensure that Ipl1-Bub1 dimerization did not affect kinetochore biorientation, we created a strain expressing Ipl1 and Ipl1-FRB both. Chromosome IV bioriented efficiently in both control and rapamycin-treated cells of this strain (Figure 2C top, Figure S2A). A kinase-dead allele of Ipl1, Ipl1^K133R^, did not cause the G2/M arrest when dimerized with Bub1 indicating that the kinase activity of Ipl1 is required for eSAC signaling^38^ (Figure 2C top). Finally, to confirm that the mitotic arrest resulting from Ipl1-Bub1 dimerization is independent of kinetochores, we used a fragment of Bub1, bub1^368-609^, comprising just the central domain, lacking the Bub3-binding ‘GLEBS’ domain and the kinase domain^39^. GFP-bub1^368-609^-2xFkbp12 did not localize to unattached kinetochore clusters in nocodazole-treated cells (Figure 2C bottom). Importantly, Ipl1 dimerization with GFP-bub1^368-609^-2xFkbp12 robustly activated the SAC.

To find sites within the Bub1 central domain phosphorylated upon its dimerization with Ipl1, we affinity purified GFP-bub1^368-609^-2xFkbp12 from rapamycin- and DMSO-treated cells (Figure S2B). In SDS-PAGE analysis, the mobility of GFP-bub1^368-609^-2xFkbp12 affinity-purified from rapamycin-treated cells was retarded, suggesting that it was post-translationally modified (Figure S2B). Mass spectrometry identified T438, S474, S475, T550, T556, and S596 in the Bub1 central domain as phosphorylated (Table S2). Importantly, these phosphorylations were significantly enriched in rapamycin-treated cells. Although our analysis identified only one of the 15 sites implicated in Bub1-Mad1 interaction, it suggests that following its rapamycin-induced dimerization, Ipl1 phosphorylates Bub1, and potentially Mad1, to drive eSAC signaling.

### Intact ABBA motif in Bub1 is essential for the Aurora B/Ipl1-mediated ectopic SAC activation

We next examined the residues in Bub1 needed for eSAC activity. Ipl1 dimerization with bub1^T453A, T455A^ did not affect cell cycle progression (Figure 2D top). Ipl1 dimerization with bub1^T453A^ (predicted to be a Cdk1 phospho-site) arrested the cell cycle, implicating Bub1(455T) as a residue critical for Ipl1-driven eSAC signaling (Figure S2C left). Bub1(485T) was also similarly found to be critical (Figure 2D top, Figure S2C and S2D). Since Ipl1 and Mps1 can both phosphorylate Bub1, we wanted to know if Mps1 becomes non-essential when Ipl1 is dimerized with Bub1. Ipl1-driven eSAC signaling was abolished when the kinase activity of an analog-sensitive Mps1 allele was conditionally inhibited^40^ (Figure S2E). Thus, Mps1 must prime Bub1 for Ipl1 activity.

We next tested whether the Bub1 ABBA motif is essential for Ipl1-driven eSAC signaling. Surprisingly, Ipl1 dimerization with bub1^-abba^ did not affect the cell cycle (Figure 2D bottom flow cytometry panel). Mps1-driven eSAC activity is unaffected by the same mutation (Figure 1C). Thus, only Ipl1-driven eSAC signaling requires the Bub1 ABBA motif to be intact. Consistently, Ipl1 dimerization with Mad1 in either cells lacking Bub1 or in cells that express either bub1^-abba^ or bub1^T485A, T509A, T518A^ did not affect the cell cycle (Figure 2E).

These data advance the following model for the direct role of Ipl1 in MCC production. Mps1 activates the SAC by sequentially phosphorylating the MELT, Bub1, and Mad1 to license the recruitment of Bub1-Bub3, Mad1-Mad2, and Cdc20, respectively. Although Ipl1 localizes to unattached kinetochores, it cannot activate the SAC on its own because it cannot phosphorylate the MELT motifs^13,14^. After Mps1 phosphorylates the MELT motifs and Bub1, Ipl1 phosphorylates Bub1 to further promote Mad1-Mad2 recruitment. Finally, Ipl1 cannot phosphorylate Mad1 to enable the Mad1-Cdc20 interaction; Mad1 must cooperate with Bub1 to promote MCC formation^32,33^.

At this juncture, it is important to note two properties of the eSAC system that likely accentuate it’s signaling activity. First, by labelling the genomic copy of Ipl1 with Frb, nearly all Ipl1 in a cell will dimerize with the Fkbp12-tagged SAC protein (depending on its relative abundance). Second, the high affinity of rapamycin-mediated dimerization of Fkbp12 and Frb will allow Ipl1 to maximally phosphorylate the signaling protein. Under physiological conditions, this Ipl1 contribution is likely to be significantly smaller^41^.

### Evidence supporting a direct role of Aurora B/Ipl1 in driving MCC assembly in kinetochore-based SAC signaling

To detect the contribution of Ipl1 to kinetochore-based SAC signaling, two conditions must prevail. First, Mps1 activity in the kinetochore must be minimal so that the Ipl1 contribution can be detected. Second, Bub1 must still be recruited to the kinetochore so that Ipl1 can act on it. As an appropriate model for these conditions, we used *spc105*^*RASA*^, a well-characterized mutant of Spc105. In this mutant, the highly conserved ‘RVSF’ motif that binds Protein Phosphatase 1 (PP1) is inactivated. Consequently, the MELT motifs remain phosphorylated, and Bub3-Bub1 remains bound to the MELT motifs in bioriented kinetochores^14^. Thus, the SAC remains active even after kinetochore biorientation^30,42,43^. The concurrence of persistent Bub1 recruitment to stably attached kinetochores with diminished Mps1 activity provides the requisite conditions for observing Ipl1 contributions to SAC signaling. Indeed, the study by Rosenberg et al. found that a hypomorphic, temperature-sensitive mutant of Ipl1 suppresses the SAC-mediated lethality of *spc105*^*RASA*^.

The Bub1 ABBA motif is necessary for Ipl1 mediated, but not for Mps1-mediated, SAC signaling. Therefore, we studied genetic interactions between *bub1*^*-abba*^ and *spc105*^*RASA*^. We found that *bub1*^*-abba*^ mutation rescued the viability of *spc105*^*RASA*^ (Figure 2F, left). We observed a similar rescue of *spc105*^*RASA*^ by *bub1*^*T455A, T485A*^ and *bub1*^*T485A, T509A, T518A*^ (Figure S2F and bub1^T453A,T455A^, also see reference^30^). Importantly, the *bub1*^*-abba*^ *spc105*^*RASA*^ mutant had a functional checkpoint (Figure 2E, bottom). These genetic interactions imply that the Bub1-mediated recruitment of both Mad1 and Cdc20 is necessary for persistent SAC signaling from bioriented kinetochores in the *spc105*^*RASA*^ mutant. Together with the requirement of Ipl1 kinase activity for SAC signaling in the *spc105*^*RASA*^ mutant, these data support our hypothesis that Ipl1 directly promotes MCC formation from unattached kinetochores.

Ipl1 may phosphoregulate the Cdc20 recruited by Bub1 to promote SAC signaling. We ruled out this possibility by studying genetic interactions between *spc105*^*RASA*^ and several Cdc20 mutants wherein the known or predicted phosphorylation sites in Cdc20 are non-phosphorylatable (data now shown).

### Aurora B drives ectopic SAC signaling in human cells

In human cells, Aurora B promotes SAC signaling indirectly by: (1) creating unattached kinetochores, (2) potentiating Mps1 recruitment to the kinetochore^10^, and (3) inhibiting the phosphatase activity antagonizing Mps1^9^. However, in fission yeast, Aurora B activity is necessary for maintaining mitotic arrest induced by unattached kinetochores^21^. Thus, the creation of unattached kinetochores is not its only role in SAC signaling. The requirement of Aurora B kinase activity for maximal SAC signaling has also been noted^20^. In human and fission yeast cells, Aurora B kinase activity and Bub1 are both required for SAC signaling driven by Mad1 that is artificially tethering to bioriented kinetochores^44-47^, similar to the requirement of Bub1 when Aurora B and Mad1 are dimerized in yeast cells (Figure 2D).

To test if Aurora B can drive eSAC signaling in HeLa cells, we adapted the previously described eSAC system^13^. We created cell lines that constitutively express mNeonGreen-2xFkbp12 fusion of either a KNL1 (Spc105 homolog) phosphodomain containing six MELT motifs (two tandem copies of a fragment spanning three motifs in KNL1^48^), the central domain of Bub1, or the C-terminal domain of Mad1 (Figure 3A, Methods). In these cells, we conditionally expressed Frb-mCherry-INCENP^818-918^. INCENP^818-918^ binds to and activates Aurora B in human cells^49^. As expected, the INCENP fragment was cytosolic, and did not show detectable localization at kinetochores (Figure S3A). We used the cell-to-cell variation in Frb-mCherry-INCENP^818-918^ expression to characterize the dependence of mitotic duration on the eSAC dosage^13^ (Figure 3A-B).

**Figure 3.**
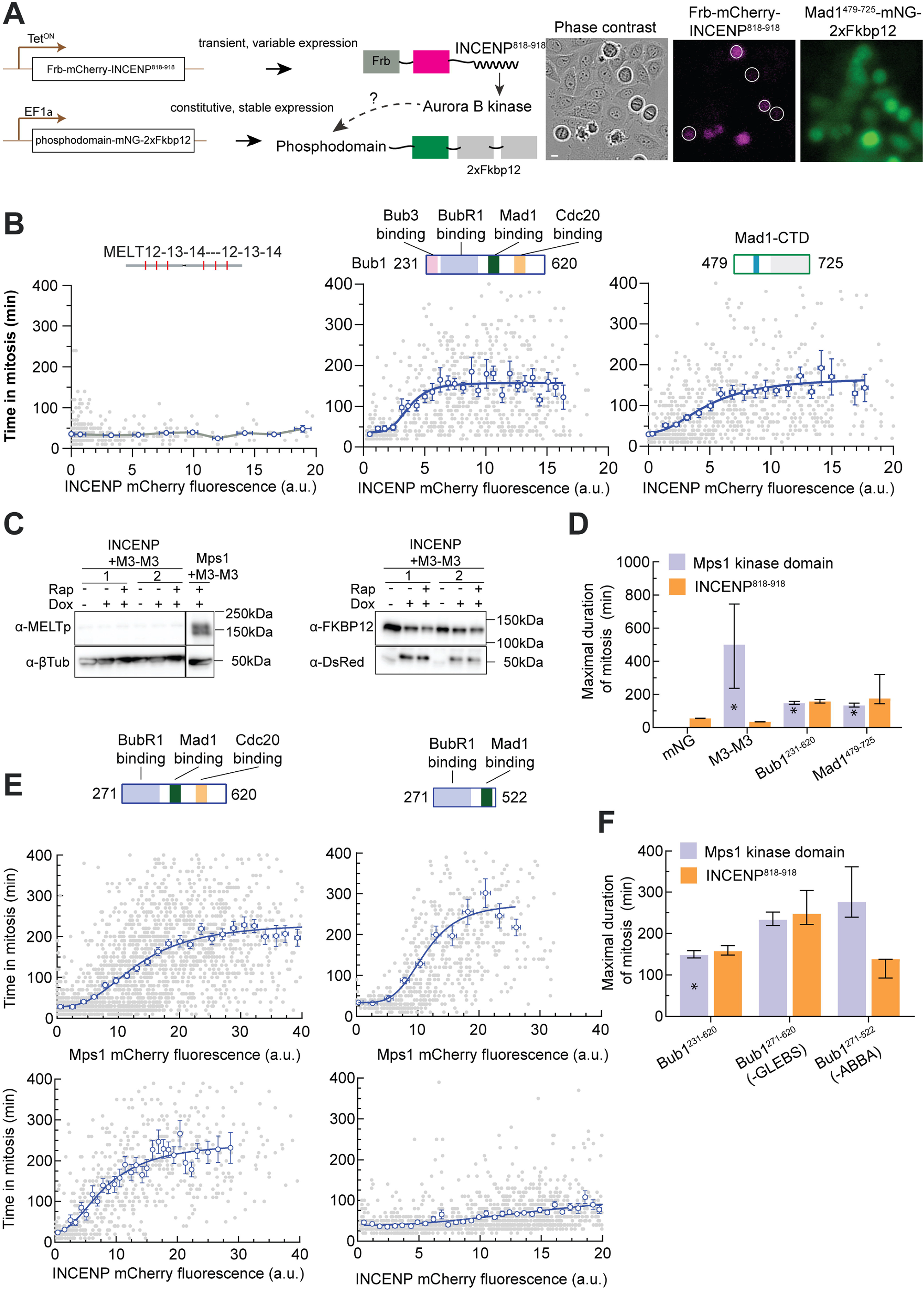
Rapamycin-induced dimerization of INCENP^818-918^ with Bub1 and Mad1, but not with MELT motifs, leads to ectopic SAC activation in HeLa cells. (A) Left: Schematic of the eSAC system designed to test the roles of Aurora B kinase activity in the core SAC signaling cascade in HeLa cells. Right: A representative micrograph from a time-lapse experiment showing the variable expression of Frb-mCherry-INCENP^818-918^. Scale bar ∼ 8.25 microns. (B) Schematic at the top displays the domain organization of the eSAC phosphodomain. Scatter plots show the correlation between time in mitosis for a given cell and the average mCherry fluorescence at the beginning of mitosis for that cell. Each gray dot represents one cell (n = 520, 787, 840 respectively, data pooled from 2 experiments). The blue circles represent the mean of data binned according to the mCherry signal; the horizontal and vertical lines represent the s.e.m. For the Bub1 and Mad1-CTD fragments, the solid blue lines display a four-parameter sigmoidal fit to the binned mean values; R^2^ values = 0.2, 0.2, respectively. (C) Western blot probing for the phosphorylation of the MEIT motif by Aurora B. Also see Figure S3C. (D) Bar graphs display the maximal response predicted by the 4-parameter sigmoidal fit from B. Vertical lines display the 95% confidence interval for the estimated maximal response. For comparison, the maximal response from eSAC systems comprised of the same three eSAC phosphodomains dimerized with the Mps1 kinase domain is also plotted (data marked by asterisks reproduced from ^13^). Vertical lines for the M3-M3-Mps1 dimerization represent the standard deviation of the bin corresponding to the peak eSAC response. This representation was made necessary by the non-monotonic nature of the dose-response data. (E) The contributions of the Bub3- and Cdc20-binding domains in Bub1 to the observed eSAC activity driven by either the Mps1 or Aurora B (n = 2635, 614, for the top panel and n = 752, 1524 for the bottom panel; R^2^ values = 0.44, 0.24, 0.51, and 0.16 respectively pooled from at least 2 experiments). The domain organization of the phosphodomain is displayed in the schematic at the top. Data presented as in B. (F) Comparison of the maximal response elicited from the indicated phosphodomains by either the Mps1 kinase domain or INCENP^818-918^ (predicted mean ± 95% confidence intervals). See also Figure S3, Data S1C-E, Video S1 and Video S2.

Rapamycin-induced dimerization of Frb-mCherry-INCENP^818-918^ with mNeonGreen-2xFkbp12 did not affect the duration of mitosis (Figure 3D), indicating that INCENP^818-918^ over-expression does not affect mitotic duration. INCENP^818-918^ dimerization with the KNL1 fragment containing six MELT motifs also did not affect mitotic progression (Figure 3B, left, Video S1 left panel). Consistently, western blot analysis of whole cell lysates following rapamycin treatment showed that the ‘MEIT’ motifs (variants of the consensus ‘MELT’ sequence) in the eSAC phosphodomain were not phosphorylated (Figure 3C, Figure S3B-C). INCENP^818-918^ dimerization with either Bub1^231-620^ or Mad1^479-725^ resulted in a dose-dependent mitotic delay (Figure 3B, middle and right, Videos S1 middle and right panels). The maximal delay in both cases was similar in magnitude, and comparable to the maximal delay caused by the dimerization of the Mps1 kinase domain with the same protein fragments (Figure 3D; data marked with an asterisk are reproduced from^13^ for comparison). Thus, Aurora B also drives eSAC signaling by phosphorylating either Bub1 or Mad1, or both.

Aurora B-driven eSAC activity persisted even when it was dimerized with a fragment of Bub1, Bub1^271-620^, lacking the Bub3-binding GLEBS motif and the N-terminal TPR domain^39,50^. Thus, the eSAC activity is kinetochore-independent (Figure 3E, also see Figure S4A). The same was true for Mps1-driven eSAC activity (Figure 3E left). In fact, the maximal delay was slightly higher and therefore the eSAC activity stronger, presumably because Bub1^271-620^ does not sequester Bub3, thus ensuring the full availability of Bub3-BubR1 for MCC formation^51^. eSAC activity also requires an active Aurora B kinase because direct dimerization of the kinase domain of Aurora B (residues 61-344), with the Bub1 phosphodomain did not affect the mitotic progression (Video S2).

Finally, we tested whether the observed mitotic delays are due to ectopic MCC formation. We synchronized HeLa cells expressing INCENP^818-918^ and Bub1^231-620^ in G2/M and then released them into media either with or without rapamycin. In each case, we harvested mitotic cells, prepared clarified cell lysates, and immunoprecipitated Cdc20. We performed the same assay for cells wherein Mps1 kinase domain is dimerized with Bub1^271-620^-mNeonGreen-2xFkbp12. In both cases, increased amounts of Mad2 co-immunoprecipitated with Cdc20 from rapamycin-treated cells compared to DMSO-treated cells (Figure S3D). Thus, INCENP^818-918^ dimerization with Bub1^271-620^ enhances Cdc20:C-Mad2 levels. The increase in Mad2 was greater in the Mps1-driven eSAC system than in the Aurora B driven system likely because Aurora B works downstream from Mps1.

### The ABBA motif of human Bub1 is necessary for the ectopic SAC activation by Aurora B

We next dimerized Bub1^271-522^, which lacks the ABBA motif, with either the Mps1 kinase domain or INCENP^818-918^. Mps1-driven eSAC activity persisted in this case, but Aurora B-driven eSAC activity was significantly weaker (Figures 3E-F). These results mirror our findings from budding yeast: Bub1 ABBA motif is dispensable for Mps1-driven eSAC activity, but necessary for Aurora B-driven eSAC activity. This observation also implies that the mitotic delay observed upon Frb-mCherry-INCENP^818-918^ dimerization with Mad1^479-725^ will require the Bub1-Mad1 interaction, which in turn requires Mps1 activity. Rapamycin-induced dimerization of INCENP^818-918^ and Mad1^479-725^ failed to delay cell division in cells treated with the Mps1-inhibitor Reversine (Figure S3E). Thus, Mps1 kinase activity is still required for the eSAC activity induced upon INCENP^818-918^ dimerization with Mad1^479-725^.

### Aurora B kinase activity promotes MCC assembly during kinetochore-based SAC signaling

To detect the contribution of Aurora B to kinetochore-based SAC signaling in HeLa cells, an additional condition must be fulfilled apart from the two mentioned earlier. Aurora B downregulates PP1 and PP2A activity unattached kinetochores, which both antagonize SAC signaling. To negate these SAC antagonists, it becomes necessary to inhibit their activity^9^ following Aurora B inhibition. To enable this, we used the following approach. We released G1/S synchronized HeLa cells expressing H2B-RFP into the cell cycle and treated them with nocodazole to activate the SAC. In these cells, we partially suppressed Mps1 kinase activity to license Bub1-Bub3 recruitment in the unattached kinetochores^48^. We suppressed PP2A recruitment to the kinetochore either by RNAi mediated knockdown of the five B56 isoforms that target PP2A to the kinetochore or by using Calyculin A^52^.

As in prior studies, partial inhibition of Mps1 significantly reduced the average duration of the mitotic arrests to 126 ± 100 minutes (mean ± S.D., Figure 4A, compared to > 1000 minutes in cells treated with nocodazole alone, data not shown). B56 RNAi increased the duration of the mitotic arrest to 258 ± 160 minutes^9^ (mean ± S.D., Video S3 left). When Aurora B activity was inhibited using the small molecule inhibitor ZM447439 under the same condition, the mitotic arrest was completely abolished (15 ± 6 minutes, Video S3 right). Calyculin A-treated cells similarly exited mitosis soon after entering it (Figure 4A). These experiments suggest that Aurora B kinase activity contributes to SAC signaling directly and independently from its indirect role in retarding SAC silencing.

**Figure 4.**
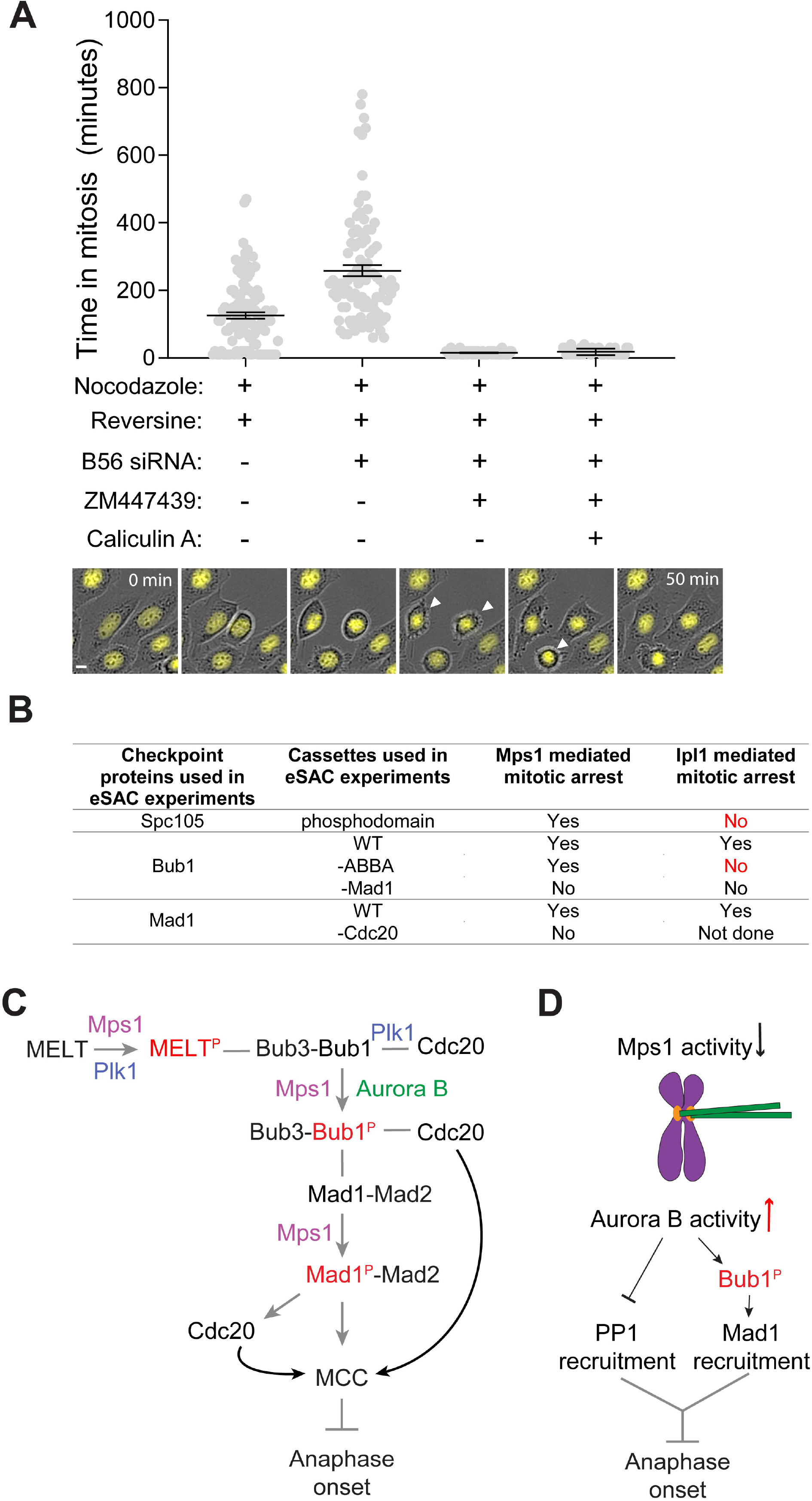
Aurora B contribution to kinetochore-based SAC signaling. (A) Scatter plot displays the duration of the mitotic arrest. Experimental treatments are indicted below each bar. (n = 92, 48, 43, 44, 41, experiment performed twice). Cells treated with B56 RNAi and ZM447439 both exited from mitosis very rapidly without assuming the rounded morphology. In this case, entry of the cell into mitosis was visually identified by the release of surface adhesion along with concurrent condensation of Histone H2B signal (in one experiment). Exit from mitosis was identified from the re-spreading off the cell over the surface (micrograph montage at the bottom, also see Video S3). Scale bar ∼ 9 microns. (B) Table summarizes the results from the eSAC experiments performed both in yeast and human cells. (C) The proposed mechanism of the direct role of Aurora B kinase activity in SAC signaling. (D) Aurora B-mediated promotion of MCC generation may enable kinetochores with syntelic attachments to continue to produce MCC and thus delay anaphase onset. See also Video S3.

In summary, our data reveal that Aurora B can phosphorylate Bub1 to promote its interaction with Mad1, and thus contribute directly to MCC formation (Figure 4B). Aurora B does not activate the SAC on its own, because it cannot phosphorylate the MELT motifs in Spc105/KNL1. After Mps1 phosphorylates the MELT motifs and Bub1^5,6^, Aurora B can act on Bub1 to promote the Mad1-Mad2 recruitment. This coordinates the interaction between Mad1-Mad2 and the Cdc20 molecule recruited by the ABBA motif in Bub1, facilitates in the formation of the closed-Mad2-Cdc20 complex and, ultimately, the MCC^32,33^. The requirement of Mps1 upstream from Aurora B in kinetochore-based SAC signaling and the significant overlap among the phosphorylation targets of the two kinases prevented us from testing this model directly. Nonetheless, it explains prior observations showing that Aurora B cooperates specifically with Bub1 in SAC signaling^18^.

Aurora B’s role in promoting MCC formation will be physiologically significant, especially when Mps1 activity in the kinetochore is weakened. One such situation occurs in kinetochores with end-on, but syntelic, attachments (Figure 4C). End-on attachments suppress Mps1 activity^53-56^ and weaken Mad1 recruitment via the SAC signaling cascade and possibly through the fibrous corona^57^. In these kinetochores, Aurora B can promote MCC formation, delay anaphase onset, and thereby reduce chromosome missegregation. Indeed, Aurora B and the ABBA motif in Bub1 are both essential for SAC signaling in Taxol-treated cells, wherein kinetochores maintain end-on attachments^17,18^. The direct role of Aurora B in SAC signaling may also contribute to the positive correlation between centromeric tension and SAC signaling.

## Acknowledgements

This work was funded by the 5R35-GM126983 from NIGMS to APJ. We thank Prof. Mara Duncan and her lab (Department of Cell and Developmental Biology, University of Michigan Medical School) for their help. We acknowledge all the Joglekar lab members for their constructive criticism. We specially thank our past lab member Alan A. Goldfarb, who did the initial pilot assay to show that rapamycin induced dimerization of Mps1-FRB with Mad1-Fkbp12 arrests the cell cycle. The authors acknowledge that this work would not have been possible without the HeLa cell line, which was developed from Henrietta Lacks’ cells taken without compensation or informed consent.

## Author Contributions

Conceptualization, Methodology, Writing – Original Draft, Writing – Review & Editing, Visualization, and Supervision, B.R. and A.P.J.; Formal Analysis and Investigation, B.R., S.J.Y.H, A.F. S.J. and A.P.J.; Software, Resources, and Funding Acquisition, A.P.J.

## STAR Methods

### Resource Availability

#### Lead Contact

Further information and requests for resources and reagents should be directed to and will be fulfilled by the Lead Contact, Ajit P. Joglekar.

#### Materials Availability

Yeast strains, HeLa cell lines and plasmids constructed in this study are available from the lead contact upon request.

#### Data and Code Availability

- Any additional information required to reanalyze the data reported in this paper is available from the lead contact upon request.

### Experimental Model and Subject Details

#### Yeast Cell culture

Yeast strains were grown in YPD (yeast extract 1%, peptone 2%, dextrose 2%) or synthetic media supplemented with 2% dextrose (as per requirement of the yeast strain) at 32°C. At the time of imaging strains with TetR-GFP and *CENIV-TetO*, we supplemented the growth media with uracil (final 40µg/ml) to distinctively observe TetR-GFP bound to *CENIV-TetO* array.

To induce meiosis or sporulation, diploid yeast strains were grown in YPD overnight to stationary phase. Next day, the cells were pelleted and resuspended them with starvation media (0.1% yeast extract, 1% potassium acetate, 0.025% dextrose) and incubated 4-5 days at RT. To obtain the strain of interest, plates with dissected tetrads were incubated for 2-3 days. Segregants with wild-type genotype or those harboring *spc105Δ, spc105*^*RASA*^, *bub1*^*- abba/phosphomutants*^ grew within that time period. Infrequently, segregants with *spc105Δ, spc105*^*RASA*^ form micro colonies after 5-6 days of incubation at 30C (Figure 2F, left). To ensure that background mutations are not responsible for the rescue of *spc105*^*RASA*^ in *bub1*^*-abba*^ background, mutant segregants were back-crossed wild-type parent strains (YEF473).

For the experiment involving analog sensitive Mps1 yeast cultures were grown for 3h till they attained mid-log phase. These cultures were treated with Hydroxyurea (100mM final) for 2h 30min to synchronize the cells late S phase. Following that, the cells were washed with YPD and released into either YPD (control) or YPD supplemented with 1-NMPP1 (50µM final). After 15min of incubation with 1-NMPP1, either DMSO (control) or rapamycin (1 µg/ml; to mediate the dimerization of Aurora B/Ipl1-Frb-GFP and Bub1-2xFkbp12) was added to the media. Mitotic cells were categorized as prometaphase, metaphase and anaphase cells according to the distribution of fluorescently labeled kinetochores within each cell (representative images in Figure S2E). The unbudded cells were considered as the cells in G1 and thus they were not taken into consideration in our analyses.

### Tissue culture and generation of stable cell lines

Henrietta Lacks (HeLa) cells were grown in DMEM media with 10% FBS, 1% Pen/Strep, and 25 mM HEPES at 37 °C and 5% CO2. Stable cell lines expressing the two eSAC components were generated by integrating a bi-cistronic eSAC plasmid at an engineered *Loxp* site in the HeLa genome according to the protocol described in ^62^. Clones with stable integration of the eSAC plasmid were selected using Puromycin (1 ug/ml), and several clones were pooled together to create the cell cultures used in the experiments.

To conduct dose-response analysis, each eSAC cell line was plated ∼ 40-48 hours prior to the start of the experiment in DMEM media without Puromycin. Doxycycline was added at the time of plating to induce the expression of either Frb-mCherry-Mps1 or Frb-mCherry-INCENP^818-918^. Prior to imaging, the cells were washed with PBS. Fluorobrite media with 10% FBS, 1% Pen/Strep with or without Rapamycin were added to each well.

### Method Details

#### Plasmid and strain construction for study involving *S. cerevisiae*

Plasmids and *S. cerevisiae* strains and cell lines used in this study are tabulated in Table S3. *S. cerevisiae* strains containing multiple genetic modifications were constructed using standard yeast genetics. Proteins tagged with GFP(S65T) and mCherry or yeast codon optimized mCherry were used to visualize kinetochores, spindle pole bodies and SAC signaling components. A 7-amino-acid peptide (sequence: ‘RIPGLIN’) was used as the linker between the proteins and their C-terminal tags (GFP, mCherry, Frb or 2xFkbp12). The cassettes for gene deletion, gene replacement and C-terminal tags were introduced at the endogenous locus through homologous recombination of PCR amplicons or using linearized plasmids ^59^. In the past, significant strain to strain variation in the intensity of mCherry-tagged kinetochore proteins or checkpoint proteins was observed. This variation arises from the inherent variability of mCherry brightness. Therefore, all Mad1-mCherry strains were created by crossing the same transformant of Mad1-mCherry (AJY1836 or AJY3741) with other strains. The deletion mutant of *NUP60* always accompanies Mad1-mCherry to disrupt Mad1localization to the nuclear envelopes ^60^. This facilitated clearer imaging and quantification of Mad1 localized to the unattached kinetochores without affecting SAC strength.

To create any diploid yeast strains, overnight cultures of a and α mating types were mixed and spotted on a YPD plate, and then incubated for approximately 3-4 hours at 32°C.

All the Spc105 mutants used in the study are chimeras of Spc105 and GFP as described previously^30,43^. Genes encoding the chimeric proteins were introduced using a cassette that consists of the 397 bp upstream and 250 bp downstream sequences of the *SPC105* open reading frame as promoter (*prSPC105*) and terminator (*trSPC105*) sequences respectively. Genes encoding GFP(S65T) at the 222^nd^ or 455^th^ amino acid positions of Spc105 were introduced by sub-cloning with an extra *Bam*HI site (Gly-Ser) upstream and *Nhe*I site (Ala-Ser) downstream from GFP. The plasmids based on pRS305 or pRS306 backbone were linearized by *Bst*EII or *Stu*I before transformations to ensure their integration at the *LEU2* or the *URA3* locus respectively.

To build *bub1* phosphomutants containing plasmids the pSK954 plasmid backbone was used^61^. pSK954 harbors the *ADH1* transcription terminator cloned within *Asc*I-*Bgl*II sites. 500bp sequence upstream of the *BUB1* start codon, 3.063kb *BUB1* ORF sequence harboring the designated mutations and 651bp *2xFKBP12* or 705bp yeast mCherry were subcloned using *Sac*II-*Asc*I sites. The ORF and *2xFKBP12* or mCherry were linked by 21bp linker which codes for RIPGILK. 350bp sequence downstream of the *BUB1* stop codon was subcloned between *Pme*I-*Apa*I to serve as the terminator. To build the strains with *bub1* phosphomutant allele, a strain hemizygous for BUB1 was used (AJY6055). The plasmids were digested by *Apa*I and *Sac*I to release 6.279kb fragment which recombined at the deleted bub1 locus replacing the *NAT1* cassette. There are two chimeras that express bub1^T453A, T455A^ (pAJ852 and pAJ896). *BUB1* ORF of pAJ852 harbors mutations of 449SR450::TG. However, upon testing no phenotypic differences were detected between the strains constructed by pAJ852 and pAJ896. Contrary to Mad1-mCherry strains, bub1-15A-mCherry strains were created via gene replacement using linear fragments (linearized with *Apa*I-*Sac*I) of pAJ923. Therefore, the difference in intensities between wild-type and bub1-15A observed in figure S1E may have been caused by differences in mCherry brightness rather than protein localization.

pSB148 which contains *IPL1* ORF flanked 1.0 Kb promoter (*prIPL1*) and 654 bp terminator (*trIPL1*) was acquired from the Biggins lab. FRB-GFP along with ADH1 terminator was first cloned within *Age*I-*Sac*I sites to generate a chimera which can express Ipl1-Frb-GFP (pAJ941). Then a 600 bp fragment. which was synthesized by Integrated DNA technologies and harbors kinase dead mutation of Ipl1 (K133R), was cloned within the *Swa*I-*Age*I sites to build pAJ940. AJY6156 (*fpr1Δ, BUB1*-1x*FKBP12*-*HIS3, DSN1*-mCherry-*HYG*) was transformed with *Stu*I digests of these plasmids where they can be integrated in *URA3* loci.

Similarly, to construct the mutants of putative phosphorylation sites of Cdc20, 506bp sequence upstream of the *CDC20* start codon and 1.833kb *CDC20* ORF sequence harboring the designated mutations were cloned into the *Sac*II-*Asc*I sites of pSK954. A *Spe*I site (ACTAGT) was inserted between the promoter and the ORF sequence. Finally, 300bp sequence downstream of the *CDC20* stop codon was cloned between *Pme*I-*KpnI* sites. To build strains with *cdc20* phosphomutant alleles, a strain hemizygous for *CDC20* was used (AJY5249). The plasmids were digested by *Kpn*I and *Sac*II to release 4.328kb fragment which recombined at the *CDC20* locus replacing the *TRP1* cassette.

### Plasmid and cell line construction for study involving HeLa cells

The plasmids used for the stable cell lines were based on the plasmids that have been described previously ^13^. Briefly, the phosphodomain was integrated into the constitutively expressed ORF of the plasmid using either *Not*I or *Asc*I and *Xho*I restriction sites. The INCENP^818-918^ fragment was integrated into the conditionally expressed ORF using *Fse*I and *Bgl*II restriction sites.

### Flow cytometry

To perform these experiments, the designated strains were grown to mid log phase. Media were then supplemented with Nocodazole (final concentration 15μg/ml) to depolymerize the spindle microtubules and activate the SAC and rapamycin (1 µg/ml) to induce the dimerization of FRB and Fkbp12 fused proteins ^63^. Samples containing approximately 0.1 OD_600_ cells were collected 0, 1, 2, 3 and 4h after nocodazole addition, and the cells were fixed using 75% ethanol, and stored at 4°C overnight. Next day, the cells washed and treated the cells with bovine pancreatic RNase (Millipore Sigma, final concentration 170ng/µl) at 37°C for 24h in RNase buffer (10mM Tris pH8.0, 15mM NaCl). Next, RNase was washed out and the cells resuspended in 1X phosphate buffered saline (pH 7.4) and stored at 4°C. These samples were incubated in Propidium Iodide (Millipore Sigma, final concentration 5µg/ml in PBS) for at least 1h at RT on the day of the assay. The stained cells were analyzed using the LSR Fortessa (BD Biosciences) in Biomedical research core facility, University of Michigan medical school. For each strain flow cytometry was performed at least twice. Representative results from one of these experiments are displayed in each panel. The data was analyzed using the FlowJO software and the graphs were adjusted by Adobe illustrator. As positive controls for SAC null phenotype, *bub1Δ* or *mad2Δ* strain were used.

### Microscopy for *S. cerevisiae* cells and image analysis

A Nikon Ti-E inverted microscope with a 1.4 NA, 100X, oil-immersion objective was used for all imaging experiments. Additionally, the 1.5X opto-var lens was used to measure Mad1-mCherry intensities. The cells were imaged at room temperature in synthetic dextrose (or synthetic galactose media whenever it was required for the assay) supplemented with essential amino acids to obtain at least 20 microscopic fields at a given time points for any strains. Mounting media were supplemented with nocodazole to image the nocodazole arrested cells. For each field of view, a ten-plane Z-stack was acquired (200nm separation between adjacent planes), and at least 20 fields were acquired in each experiment.

Total fluorescence intensities of kinetochore clusters (16 kinetochores in metaphase) were measured by integrating the intensities over a 6×6 region centered on the maximum intensity pixel. Median intensity of pixels immediately surrounding or a nearby 6×6 area was used to correct for background fluorescence. Fluorescence intensity was calculated as described previously ^64,65^.

### Drug treatments and RNAi for experiments with human cells

To induce the expression of either mCherry-Frb-Mps1 kinase domain or -Incenp^818-918^, doxycycline was added to a final concentration of 2 ug/ml (stock concentration 2 mg/ml in DMSO). To induce the dimerization of protein fragments, Rapamycin was added ∼ 1 hour prior to the start of the experiment to a final concentration of 500 nM (stock concentration 500 μM in DMSO). GSK-923295 was added to the final concentration of 15 nM (stock concentration 236 µM). Partial Mps1 inhibition was achieved by adding Reversine to the final concentration of 250 nM (stock concentration 500 μM in DMSO). Nocodazole was added to the final concentration of 330 nM (stock concentration 330 μM in DMSO). ZM447439 was added to the final concentration of 10 μM (stock concentration 3mM in DMSO). Calyculin A was added to the final concentration of 100nM (stock concentration 50µM in DMSO). The cocktail of siRNA against five different B56 isoforms was added to a final concentration of 40 nM (stock concentration 10 μM). The siRNA sequences were obtained from ref. ^9^.

### Long term live Cell Imaging of HeLa cells and Image analysis

Imaging was conducted over a period of 24 hours as described in detail previously ^13^. Either the Incucyte Zoom Live Cell Imaging system (Sartorus Inc.) or the ImageExpress Nano live cell imaging system (Molecular Devices) using 20x Phase objectives for imaging. To image cells on the Incucyte system, cells were plated in 12-well plastic tissue culture plates, whereas they were plated in 24-well plate glass-bottom dishes for imaging using the ImageExpress Nano system. Typically, 4 positions were selected within each well for imaging. At each position, one phase, GFP, and mCherry image was acquired every 10 minutes. The exposure time for mCherry image was adjusted to minimize photobleaching while ensuring accurate determination of cellular intensity values. It should be noted that the excitation intensity of the Incucyte instrument declined significantly over the course of this study. Furthermore, a small minority of the experiments were carried out on the ImageExpress Nano microscope, which has excitation sources, optics, and detector that are entirely different from the components of the Incucyte microscope. Therefore, the mCherry intensity values across different experiments are not directly comparable. The duration of mitosis and GFP and mCherry fluorescence per cell were determined using a custom image analysis script implemented by a Matlab graphical user interface as described previously ^13^.

### Immunofluorescence assay in HeLa cells

Immunofluorescence assay was performed as described previously (Deluca et al., 2011, J Cell Sci). We plated cells expressing Frb-mCherry-INCENP^818-918^ on a 12mm coverslip until they reached 80% confluence. The cells were treated with 15 nM GSK-923295 for 3 h to activate SAC. During fixation, the cells were pre-extracted with 0.5% Triton X-100 in 0.1 M PHEM (240 mM Pipes, 100 mM HEPES, 8 mM MgCl2 and 40 mM EGTA) and fixed with 4% PFA for 10 minutes. The coverslips were washed three times with 0.1 M PHEM and blocked for 30 min at room-temperature with StartingBlock (Thermo scientific, 37578). Next, the cover slips were incubated in primary antibody (Mouse anti-Hec1, 1:3000) over night at 4□ C. Next day, the coverslips were washed 4 times with 1xPHEM containing 0.05% Tween-20 and incubated with secondary antibodies (Goat anti-Mouse 488, 1: 10000) for an additional 45 min at room temperature in dark. Following that, the coverslips washed again four times and mounted in antifade (ProLong; Molecular Probes).

Immunofluorescence images were taken on a Nikon inverted microscope equipped with a Crest X-Light V2 LFOV25 Spinning Disk Confocal head, and Photometric 95B Prime cMOS camera. For each field of view, 31 Z sections were acquired at 0.2 mm steps using a 100x 3, 1.4 Numerical Aperture (NA) Nikon objective.

### Immunoprecipitation of Cdc20

This assay was performed as described in ^66^. Cells expressing Bub1^271-620^-neon2xFkbp12, Frb-mCherry-Mps1 or Bub1^225-620^-mNeon-2xFkbp12, Frb-mCherry-Incenp^818-918^ were grown to ∼ 50% confluence. These cells were synchronized in S phase using double thymidine treatment (final concentration 2.5 mM). At the same time, doxycycline (final concentration 2 ug/ml) was added to the media to induce expression of Frb-mCherry-Mps1 kinase domain or Frb-mCherry-INCENP^818-918^. G1/S synchronized cells were washed with PBS and released into media supplemented with the Cdk1 inhibitor RO-3306 (final concentration 15 nM). After 20 hours, G2-synchronized cells were washed with PBS and released into media supplemented with rapamycin (1 µg/ml) for 4 h.

Mitotic cells were harvested using plate shake-off followed by centrifugation at room temperature/ 1100 rpm/ 5 min. Cells were treated with a volume of lysis buffer (75 mM HEPES, pH 7.5; 150 mM KCl, 1.5 mM MgCl_2_, 1.5 mM EGTA, 10% glycerol, 1% CHAPS, supplemented with protease inhibitor and phosphatase inhibitor cocktails) added in proportion to the weight of the cell pellet. After a 10 min incubation on ice, total cell lysates were clarified by centrifugation (4□C/ 13,000 rpm for 10 min). The cell lysates were incubated with α-Cdc20 anitbody-conjugated protein G-Sepharose 4B beads (4 µg antibodies per 10 µl of beads, Thermo Scientific, catalog #20399) at room temperature for at least 1 h. After that, the beads were washed three times with lysis buffer and boiled in Laemmli sample buffer.

### Affinity purification of Bub1 middle domain and mass spectrometry

Mid-log phase cells expressing Ipl1-Frb and GFP-Bub1^368-609^-2xFkbp12 were treated with either DMSO (control) or Rapamycin for approximately 3 h. GFP-Bub1^368-609^-2xFkbp12 was immunoprecipitated from clarified cell lysates using GFP-TRAP beads following the methodology described previously^43,67^. Precipitated proteins were eluted into Laemmli sample buffer and subjected to SDS-PAGE analysis (See Figure S2B). Incisions ∼0.5cm tall were made in the Coomassie stained SDS-PAGE, and bands were further processed for tandem mass spectrometry at the University of Michigan proteomics resource facility (See Figure S2B). Mass spectrometry data was filtered to retain peptides/peptide spectrum matches (PSM) with false discovery rate (FDR) <1%. The detected sites and flanking sequences are included in the table S2. The fold change from DMSO to rapamycin was calculated using normalized peptide abundance.

### Immunoblotting

Western blotting was performed using commercial antibodies. The specifications of antibodies and their dilutions are as follows: Mouse α-GFP, 1: 3,000; mouse α-Ds-Red, 1: 2,000; mouse α-βTubulin, 1: 15,000; rabbit α-Fkbp12, 1: 5,000; rabbit α-PhosphoMELT (MEIpT, MELT13/17), 1:2000; mouse α-Cdc20, 1:500; mouse α-Mad2, 1:1000; mouse α-BubR1, 1:1000; mouse α-Apc3, 1:1000. The primary antibodies were detected using HRP conjugated secondary antibodies (1: 10,000) per the manufacturer’s instructions. The subsequent chemiluminescence was detected using the C600 imager from Azure Biosystems. The band intensities were measured by ImageJ^68^.

### Molecular weights of the target proteins

Molecular weights of the proteins targeted in the immunoblots and Coomassie stained gels in this study, obtained from Saccharomyces Genome Database and Uniprot: yeast α-Tubulin-49.8 kDa, yeast GFP-Mad1-115.2 kDa, yeast GFP-Mad3-87 kDa, yeast GFP-bub1^368-609^-2xFkbp12-79.2 kDa, human Frb-mCherry-INCENP^818-918^-50 kDa, human β-Tubulin-50 kDa, M6-mNeonGreen-2xFkbp12-101.0 kDa, human Mps1 (Kinase)-Frb-mCherry-80.0 kDa, human Mad2-23.5 kDa, human Cdc20-54.7 kDa, human BubR1-119.54 kDa, human Apc3-91.86 kDa.

### Quantification and Statistical Analysis

The technical replicates represent the number of times each experiment was performed. The biological replicates are defined as multiple transformants or segregants of the same strain which contain identical genotype. For imaging experiments, the number of cells analyzed for each strain and number of experimental replications is noted in the figure legends. All statistical analysis was performed using GraphPad Prism (version 8). The data were normalized with the mean intensities obtained for wild-type controls in each experiment to prepare the scatter plots of Mad1 intensities. To compare sample means in all other cases, either the t-test or two-way ANOVA test was applied to ascertain the statistical significance of the rest of the data using GraphPad Prism (version 8). The p-values obtained from these tests are indicated in the figures.

**Data S1. Complete images of the immunoblot membranes and gel pictures that were used in this study. Related to Figure S1G, S2B, 3C, S3A-C**. (A) Complete images of the immunoblot membranes probed with anti-αTub and anti-GFP in Figure S1G. Strain details of the cell lysates which were run on the gel are indicated at the top. On the left, the numbers depict the bands of molecular weight marker that was ran along with the lysates. (B) Image of the Coomassie stained whole gel where elutes from GFP-Trap beads were run. Related to Figure S2B. On the left, molecular weight markers are mentioned. (C) Images of the entire immunoblots where lysates of the indicated cell lines were probed with combination of anti-phosphoMELT and anti-βTub (left) or with that of anti-DsRed and anti-Fkbp12 (right). Related to Figure 3C and Figure S3B. Antibody dilutions used in the immunoblots are depicted at the bottom. The numbers on the side of the blots indicate the molecular weight markers. Precise molecular weights of the targeted proteins are also mentioned at the bottom. (D) Images of the complete immunoblots where lysates of the indicated cell lines were probed with combination of anti-phosphoMELT and anti-βTub (left) or with that of anti-DsRed and anti-Fkbp12 (right) in Figure S3A. Antibody dilutions used in the immunoblots are depicted at the bottom. The numbers on the side of the blots state the molecular weight markers. Precise molecular weights of the targeted proteins are also stated at the bottom. (E) Images of the immunoblot membranes where input and immunoprecipitated samples after anti-Cdc20 immunoprecipitation experiments were run in Figure S3C. The membrane was cut into three to four strips that were then probed with anti-Cdc20, anti-Mad2, anti-BubR1 and anti-Apc3 antibodies. Precise molecular weights of the targeted proteins are listed on the right of Mps1-eSAC assay immunoblots. The assay was replicated at least thrice for proper quantification and statistical significance.

## Supplementary Videos Legend

**Video S1– Effect of rapamycin induced dimerization of Frb-mCherry-INCENP**^**818-918**^ **and M3-M3-mNeonGreen-2xFkbp12 (left) or Bub1**^**231-620**^**-mNeonGreen-2xFkbp12 (middle) or Mad1**^**479-725**^**-mNeonGreen-2xFkbp12 (right) on the duration of mitosis in HeLa A12 cells (hh:mm)**. Images acquired on the Incucyte microscope. **Related to Figure 3B**.

**Video S2 – Effect of rapamycin induced dimerization of Frb-mCherry-aurora B**^**61-344**^ **with Bub1**^**231-620**^**-mNeonGreen-2xFkbp12 on the duration of mitosis in HeLa A12 cells (hh:mm)**. Images acquired on the Incucyte microscope. **Related to Figure 3B**.

**Video S3 – Left: Cell cycle progression of H2B-RFP expressing HeLa A12 cells treated with 330 nM nocodazole, 250 nM Reversine, and a cocktail of siRNA against five B56 isoforms (hh:mm). Right: Cell cycle progression of H2B-RFP expressing HeLa A12 cells treated as in left along with 10** μ**M ZM447439 (hh:mm)**. Images acquired on the ImageExpress Nano microscope. **Related to Figure 4A**.

**Table S3. list of *S. cerevisiae* strains and list of plasmids constructed and used in this study**. List of yeast strain is related to Figure 1, 2, S1 and S2. Plasmid list is related to Figure 1-4 as well as Figure S1-S3. **Related to Star Method and key resource table**.

