## Supplementary material for "Aurora B phosphorylates Bub1 to promote spindle assembly checkpoint signaling": Document S1

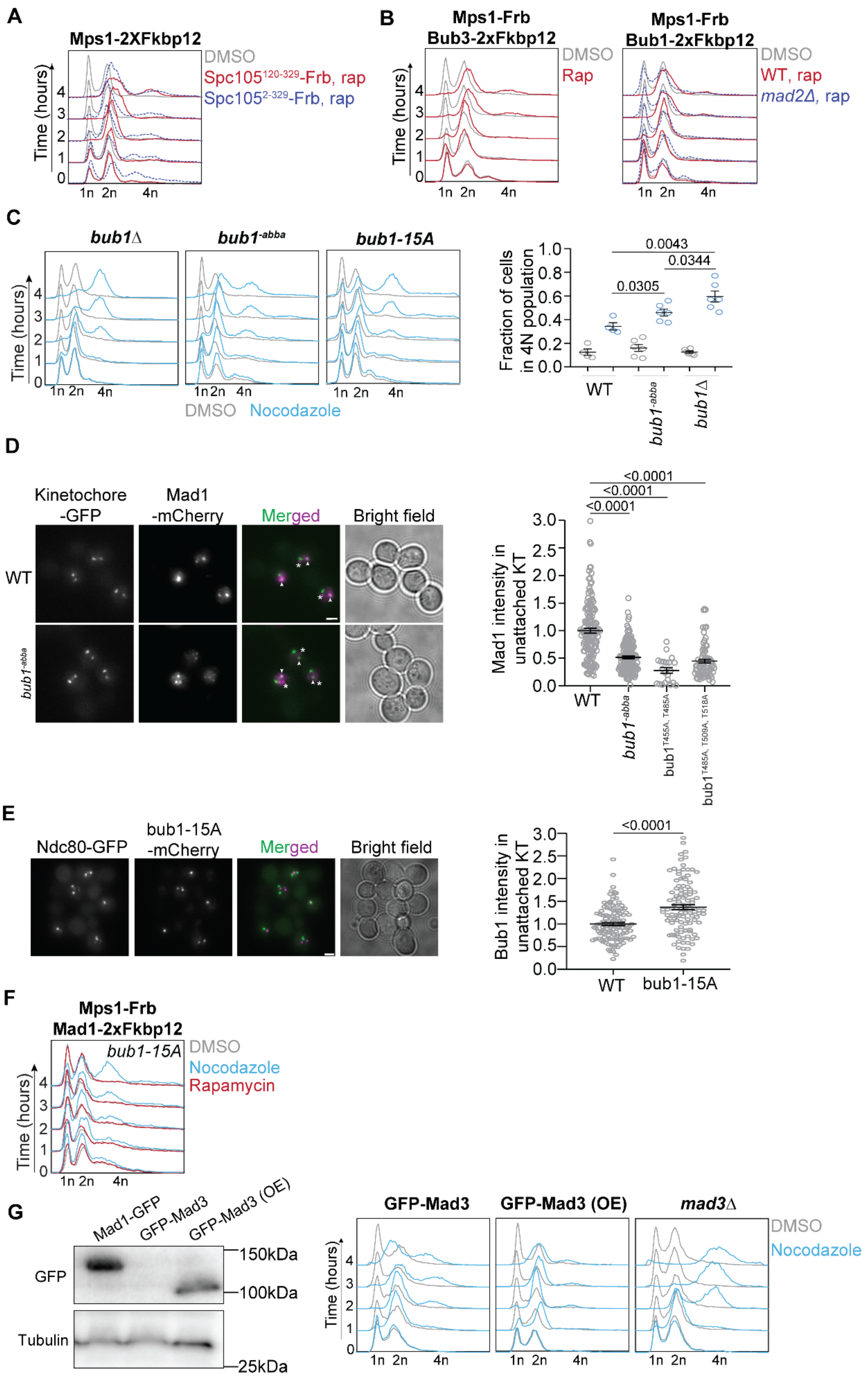
A
Mps1-2XFkbp12
DMSO
Spc105120-329-Frb, rap
Spc1052-329-Frb, rap
Time (hours)
1n 2n 4n
B
Mps1-Frb
Bub3-2xFkbp12
DMSO
Rap
Mps1-Frb
Bub1-2xFkbp12
WT, rap
mad2Δ, rap
C
bub1Δ
bub1-abba
bub1-15A
DMSO Nocodazole
Fraction of cells in 4N population
0.0043
0.0344
0.0305
WT
D
Kinetochore -GFP
Mad1 -mCherry
Merged
Bright field
Mad1 intensity in unattached KT
<0.0001
bub1T455A, T485A
bub1T485A, T509A, T518A
E
Ndc80-GFP
bub1-15A -mCherry
Bub1 intensity in unattached KT
F
Mps1-Frb
Mad1-2xFkbp12
Rapamycin
G
Mad1-GFP
GFP-Mad3
GFP-Mad3 (OE)
GFP
Tubulin
150kDa
100kDa
25kDa
mad3Δ

**Figure S1- Delineating the roles of Mps1 in the SAC signaling cascade using the 'eSAC' system. Related to Figure 1. (A)** Flow cytometry-based analysis of cell cycle progression following rapamycin-induced dimerization of Mps1-Fkbp12 with either GFP-Spc105$^{120\text{-}329}$-Frb (red), which contains six MELT repeats, or GFP-Spc105$^{2\text{-}329}$-Frb (dashed blue), which contains the Glc7 recruitment domain along with the six MELT repeats. **(B)** Left: Flow cytometric analysis of cell cycle progression following the rapamycin-induced dimerization of Bub3 and Mps1 in wild-type cells. Right: Flow cytometry analysis following the rapamycin-induced dimerization of Bub1 and Mps1 in wild-type (red) and *mad2Δ* cells (dashed blue). (**C**) Left: Effect of nocodazole treatment on cell cycle progression in *bub1Δ* (left), bub1$^{-abba}$ (2nd from the left) and bub1-15A (2nd from the right) cells. Right: Scatter plot showing quantification of fraction of 4N population in WT, bub1$^{-abba}$ and *bub1Δ* cells. The p values derived by pairwise t-tests performed on the data are mentioned on the top of the graph. **(D)** Left: Representative images of cells with unattached kinetochore clusters showing co-localization of the indicated proteins with fluorescently labeled kinetochores (Spc105$^{222::GFP}$ and Ndc80-GFP for WT and bub1$^{-abba}$ respectively). Note that the Mad1-mCherry puncta marked with an asterisk result from the deletion of the nuclear pore protein Nup60. These puncta are not associated with kinetochores. Mad1 puncta co-localizing with kinetochores are marked with arrowheads. Scale bar ~3.2µm. Right: Scatter plot shows the quantification of fluorescence signal of kinetochore-localized Mad1-mCherry (mean±s.e.m., normalized to the average signal measured in nocodazole-treated wild-type cells). n=146, 151, 21 and 73 for WT, bub1$^{-abba}$, bub1$^{T455A,\ T485A}$ and bub1$^{T485A,\ T509A,\ T518A}$ respectively pooled from two different technical repeats. The p values obtained from pairwise t-test analyses performed on the data are mentioned in the top of the plot. **(E)** Left: Representative microscopic images showing localization of bub1-15A-mCherry and unattached kinetochores (visualized by Ndc80-GFP) in yeast cells arrested in mitosis due to nocodazole treatment. Scale bar ~3.2µm. Right: Scatter plot shows the quantification of fluorescence signal of bub1-15A-mCherry (mean+s.e.m., normalized to the average signal measured in nocodazole-treated wild-type cells). n=115 and 108 for WT and bub1-15A respectively pooled from two different technical repeats. We performed Mann Whitney test on the data. The p-value is depicted above the plot. (**F**) Effect of nocodazole, rapamycin and DMSO (control) treatment on cell cycle progression in strain expressing Mps1-Frb, Mad1-2xFkbp12 and bub1-15A-mCherry. **(G)** Left: Western analysis by α-GFP and α-Tubulin (loading control) on the lysate of Mad1-GFP, GFP-Mad3 and cells wherein GFP-Mad3 is overexpressed. Right: Flow cytometry to test the effect of nocodazole treatment on either GFP-Mad3 (left), GFP-Mad3 overexpression (middle) or *mad3Δ* cells (right). Also see Data S1A.

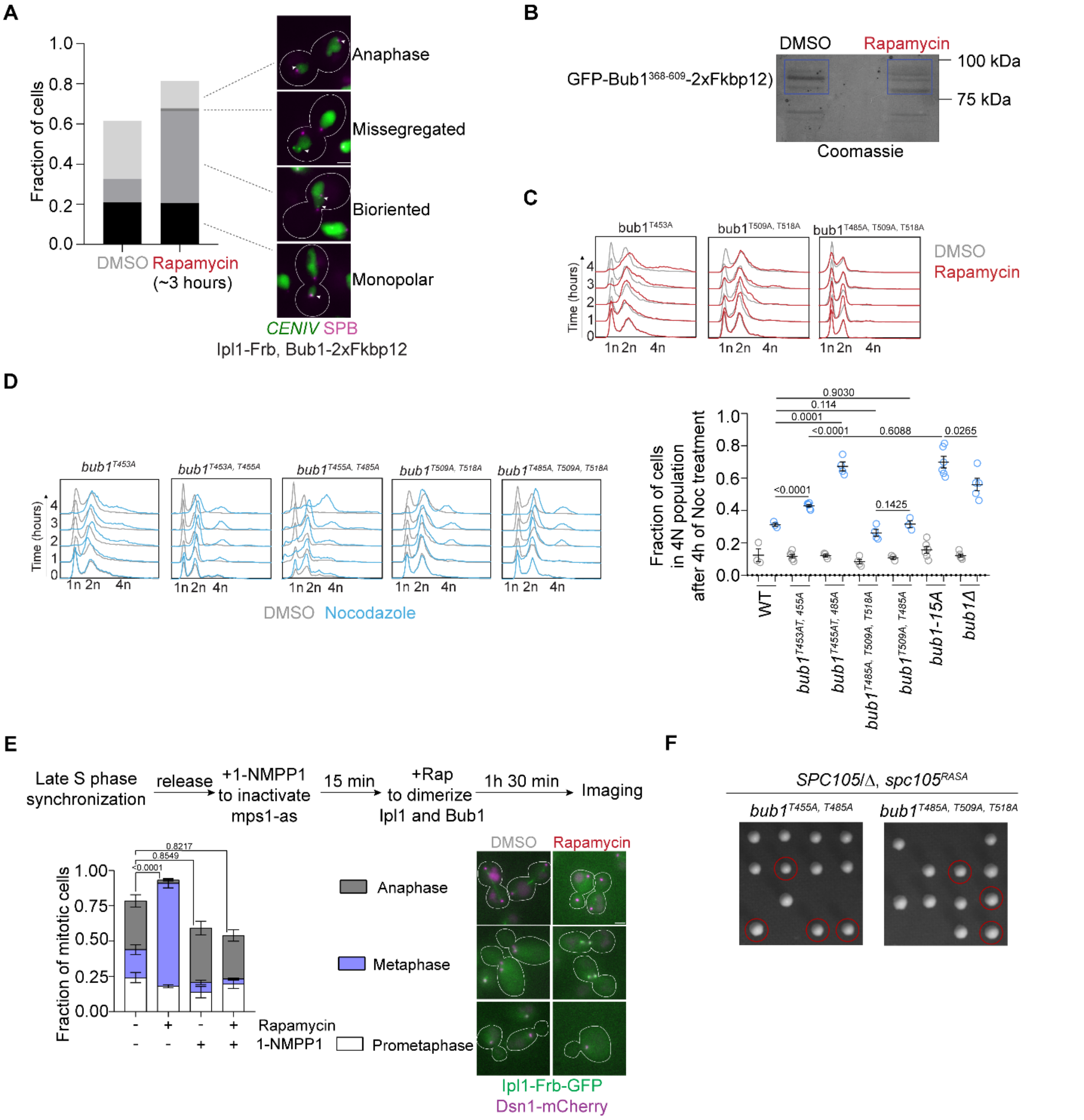


**Figure S2- Dissection of the contributions of Aurora B/Ipl1 kinase activity to SAC signaling in budding yeast, related to Figure 2. (A)** Left: Bar graph summarizes the scoring of cells based on chromosome 4 segregation (observed by TetR-GFP bound to TetO array adjacent to centromere 4/*CENIV*) and spindle morphology (Spc98-mCherry). The scoring key is displayed on the right. The number of cells scored for the analysis: n = 110 and 282 for control and rapamycin-treated cell samples respectively, pooled from three technical repeats performed on two biological replicates). Scale bar in the inset images: ~3.2 μm. Arrowheads depict the location of *CENIV-TetO* foci. **(B)** GFP-Trap purification of GFP-bub1$^{368\text{-}609}$-2xFkbp12. Image of a Coomassie stained gel following the affinity purification. Expected molecular weight of GFP- bub1$^{368-609}$-2xFkbp12 is approximately 79.0 kDa. The blue rectangles indicate the region incised and processed for mass spectrometry. Also check Data S1B for complete Coomassie stained gel image. **(C)** Effect of rapamycin induced dimerization of Ipl1 with either bub1$^{T453A}$ (left) or bub1$^{T509A,\ T518A}$ (middle) or $^{bub1T485A,\ T509A,\ T518A}$ (right). **(D)** Left: Flow cytometry profile showing cell cycle progression in nocodazole treated cells of indicated strains. Cells expressing bub1$^{T455A,\ T485A}$ transition to 4n ploidy after 4 hours of exposure to nocodazole, suggesting that the mutation weakens, but does not abolish, the SAC (compare with results of a similar analysis on *bub1Δ* cells in Figure S1D). Right: Scatter plot depicting the fraction of cells with 4N ploidy after 4 hours of nocodazole treatment (genotypes indicated below the X axis). The data were accumulated from at least three independent flow cytometry experiments. The statistical significances were derived by pairwise t-tests and p values are mentioned at the top. (**E**) Top: Flow chart describes the workflow to inactivate Mps1 prior to dimerize Ipl1-Frb-GFP with Bub1-1xFkbp12. Bottom left: Bar graph depicts the fraction of mitotic when the cells expressing analogue sensitive Mps1 were treated with rapamycin and 1-NMPP1 as mentioned in the graph. Bottom right: representative images of prometaphase, metaphase and anaphase cells as observed in untreated (DMSO) and rapamycin treated cells. The number of cells analyzed in each of these 4 treatments: n=366, 461, 246 and 974 respectively. The whole experiment was replicated four times. The statistical significances were derived by 2-way ANOVA test and p values for fractions of anaphase cells are mentioned at the top of the graph. Scale bar ~3.2µm. (**F**) Tetrad dissection analysis of the indicated strains. Please see method section for detailed description of sporulation, isolation of tetrads and obtaining segregants.

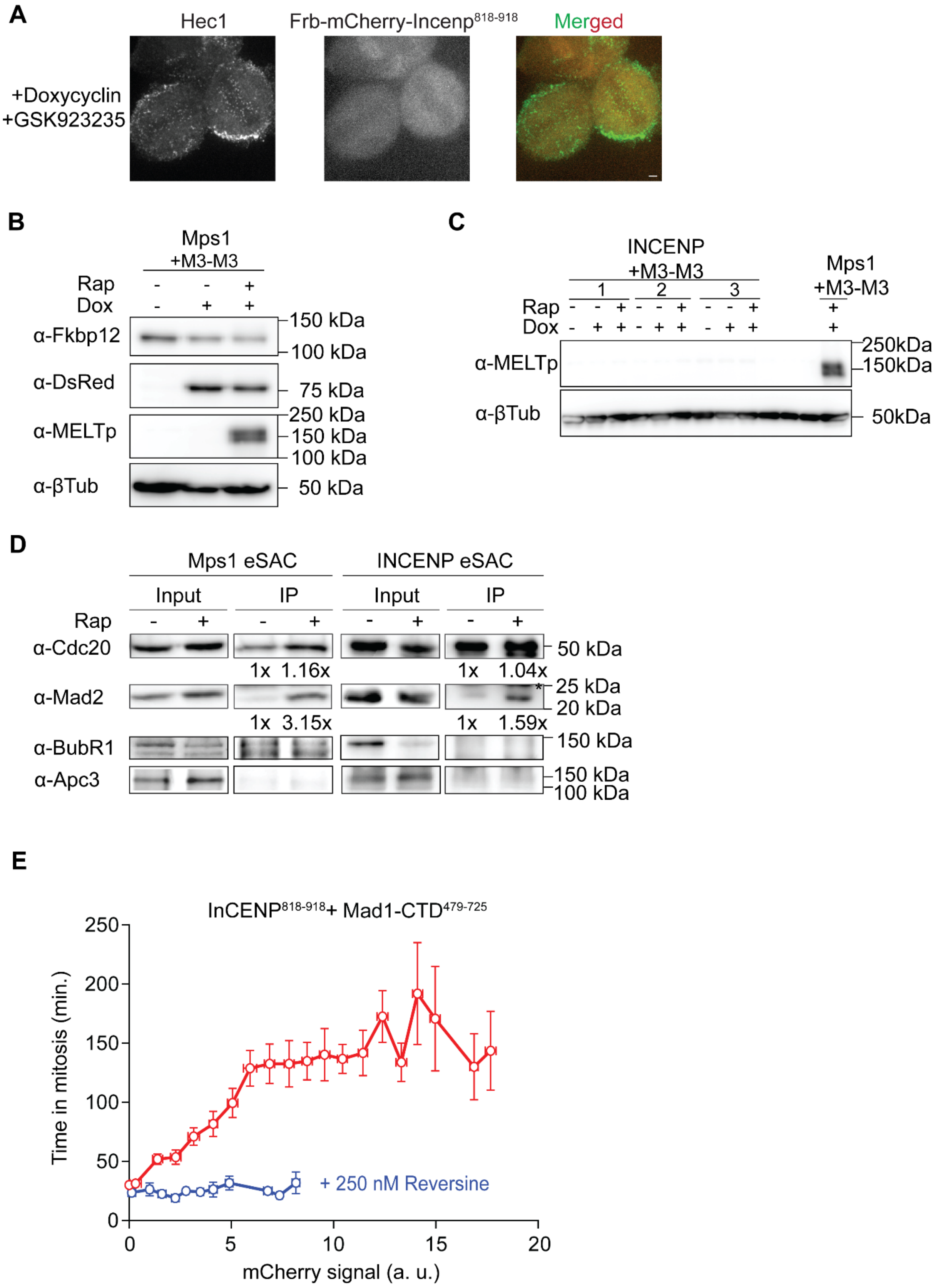

A
Hec1
Frb-mCherry-Incenp818-918
Merged
+Doxycyclin
+GSK923235
B
Mps1
+M3-M3
Rap
Dox
α-Fkbp12
α-DsRed
α-MELTp
α-βTub
150 kDa
100 kDa
75 kDa
250 kDa
150 kDa
100 kDa
50 kDa
C
INCENP
+M3-M3
1
2
3
Mps1
+M3-M3
Rap
Dox
α-MELTp
α-βTub
250kDa
150kDa
50kDa
D
Mps1 eSAC
INCENP eSAC
Input
IP
Rap
α-Cdc20
α-Mad2
α-BubR1
α-Apc3
1x 1.16x
1x 1.04x
1x 3.15x
1x 1.59x
50 kDa
25 kDa
20 kDa
150 kDa
150 kDa
100 kDa
E
InCENP818-918+ Mad1-CTD479-725
Time in mitosis (min.)
mCherry signal (a. u.)
+ 250 nM Reversine
0
50
100
150
200
250
5
10
15
20

**Figure S3 – Investigating the roles of Aurora B in SAC signaling in HeLa cells. Related to Figure 3. (A)** Representative confocal image shows localization of Hec1 (visualized by anti-Hec1$^{9G3}$ staining) and Frb-mCherry-INCENP$^{818\text{-}918}$. Cells were treated with 15 nM GSK923295, a small-molecular inhibitor of the mitotic kinesin CENP-E, for 3 h before fixation to enrich the mitotic population. Scale bar~1.462 µm. **(B)** Control western blot analysis for the phosphorylation of the MEIT motif by Mps1**.** The assay was repeated twice. Molecular weight markers are mentioned on the right. Also check Data S1D. **(C)** Immunoblot assay to analyze phosphorylation of MEIT motif by Aurora B (Frb-mCherry-INCENP$^{818\text{-}918}$). Three biological replicates were run in this gel. Here we treated the cells as indicated in the figure. As a control, we ran the lysate of doxycycline and rapamycin treated cells that express Frb-mCherry-Mps1 kinase and M3-M3-mNeongreen-2xFkbp12. Top blot was probed with α-MEITp antibodies and bottom one was probed with α-βTubulin antibodies. The molecular weight markers are mentioned on the right. See also Data S1C for immunoblot images. **(D)** Co-immunoprecipitation of Mad2, BubR1 and Apc3 with Cdc20 in Hela cell lines where we induced rapamycin mediated dimerization of Mps1 and Bub1 or Aurora B and Bub1. Numbers below Cdc20 and Mad2 lanes indicate the mean band intensities relative to that of DMSO treated samples (-rap control). Mad2 intensities were also normalized by Cdc20 band intensities. The experiment was replicated thrice. Mad2 (molecular weight 23.5 kDa, according to Uniprot) runs close to IgG light chain (marked with asterisk. In samples involving Frb-mCherry-Mps1, we observed an extra band for BubR1 that we did not see in samples with Frb-mCherry-INCENP$^{818\text{-}918}$. See also Data S1E for uncropped immunoblot images. **(E)** Partial inhibition of Mps1 due to Reversine treatment (250 nM, blue line) abolishes eSAC activity induced by the dimerization of INCENP$^{818\text{-}918}$ with Mad1$^{479\text{-}725}$(n = 154, experiment performed once). Red points and line show DMSO treated cells (data re-plotted from Figure 3B).

| Bub1 mutant | Kinase | Mitotic arrest in nocodazole | Mitotic arrest in rapamycin |
|---|---|---|---|
| T453A | Mps1 | yes | yes |
| | Ipl1 | yes | yes |
| T453A, T455A | Mps1 | yes | yes |
| | Ipl1 | yes | **No** |
| T455A, T485A | Mps1 | **No** | yes |
| | Ipl1 | **No** | **No** |
| T485A, T509A, T518A | Mps1 | yes | yes |
| | Ipl1 | yes | **No** |
| T509A, T518A | Mps1 | yes | yes |
| | Ipl1 | yes | yes |
| 15A | Mps1 | **No** | **No** |
| | Ipl1 | **No** | **No** |
| -abba | Mps1 | yes | yes |
| | Ipl1 | yes | **No** |

**Table S1. Our observations obtained from the eSAC experiments performed in yeast. Related to Figures 1,2, S1 and S2.**

| Sites in Bub1 | Peptide sequence | Fold change in rapamycin w.r.t. DMSO control |
|---|---|---|
| T438 | NLAHE**T**PVKPS | 37.31 |
| S474 | FNQHY**S**TPGAL | 5.03 |
| T475 | QHYS**T**PGALL | 2.97 |
| T550 | KADYM**T**PIKET | 9.13 |
| **T566** | VPIIQ**T**PKEQI | 4.49 |
| S596 | TTIQS**S**PFLTQ | 14.96 |

**Table S2. Phosphorylated sites on Bub1 middle domain identified by mass-spectrometry. Related to Figure 2 and S2B.** Mass spectrometry was performed on the elutes of GFP-trap assay done with the lysates of DMSO or rapamycin treated cells of fpr1Δ, IPL1-FRB, GFP-bub1368-609-2xFKBP12 (please check methods section for details of GFP-trap assay and mass spectrometry). The data was analyzed with GFP-bub368-608-2xFkbp12 sequence. The GFP-Trap assay followed by mass-spectrometry was repeated twice. The site of T566 (marked in bold red) was discovered previously as one of 15 phosphorylation sites of Bub1 middle domain [S1]. The abundance of each phosphopeptide was divided by the abundance of GFP-bub1368-609-2xFkbp12 in DMSO sample to calculate the fold-change in the rapamycin sample over the DMSO sample.
