## Supplementary material for "Aurora B phosphorylates Bub1 to promote spindle assembly checkpoint signaling": Table S3

| Strain (AJY#) | Background | Genotype description | Used in the figure |
| --- | --- | --- | --- |
| 5452 | BY4741 | *MAT a, fpr1Δ, MPS1-FRB:KAN, BUB1-2xFKBP12-HYG, SPC25-mCherry-HYG* | S1B |
| 5464 | BY4741 | *MAT a, fpr1Δ, MPS1-FRB-KAN, BUB1-2xFKBP12-HYG, mad2Δ::NAT* | S1B |
| 5364 | YEF473 | *MAT a, fpr1Δ, bub1Δ::TRP1::bub1-abba^(F490A,V492A,F493A,N495A)^-2XFKBP12(KAN), MPS1-FRB-NAT* | 1C |
| 5754 | YEF473 | *MAT a, fpr1Δ, bub1Δ::TRP1::bub1-15A-2XFKBP12 (KAN), MPS1-FRB-NAT* | 1C, S1D, S2G |
| 3536 | YEF473 | *MAT a, fpr1Δ, MAD1-2xFKBP12-HYG, MPS1-FRB-KAN* | 1D |
| 4183 | BY4741 | *MAT a, fpr1Δ, spc105Δ::NAT, MAD1-2xFKBP12-HYG, MPS1-FRB-KAN, spc105-6A (LEU2)* | 1D |
| 4184 | BY4741 | *MAT a, fpr1Δ, spc105Δ::NAT, MAD1-2xFKBP12-HYG, MPS1-FRB-KAN, spc105-6A (LEU2)* | 1D |
| 4990 | BY4741 | *MAT a, fpr1Δ, MAD1-2xFKBP12-HYG, MPS1-FRB-KAN, mad3Δ::lox-LEU2-lox* | 1D |
| 6136 | YEF473 | *MAT a, fpr1Δ, bub1Δ::TRP1::*bub1-abba^(F490A,V492A,F493A,N495A)^*-ymCherry (KAN), MAD1-2xFKBP12-HYG, MPS1-FRB-NAT* | 1D |
| 6137 | YEF473 | *MAT a, fpr1Δ, bub1Δ::TRP1::*bub1-abba^(F490A,V492A,F493A,N495A)^*-ymCherry (KAN), MAD1-2xFKBP12-HYG, MPS1-FRB-NAT* | 1D, S1D |
| 4816 | BY4741 | *MAT a,* *fpr1Δ,* *MAD1-2XFKBP12-HYG, MPS1-FRB::KAN*, *bub3Δ::NAT* | 1D |
| 5236 | BY4741 | *MAT a, fpr1Δ, MPS1-FRB-KAN, GFP-mad1^CTD^-2xFKBP12-HIS3* | 1D |
| 5517 | BY4741 | *MAT a, fpr1Δ, MPS1-FRB-KAN, mad1Δ::NAT, GFP-mad1^CTD-4A(T624A+T704A+T737A+T739A)^-FKBP12-HIS3* | 1D |
| 5142 | BY4742 | *MAT α, mad3Δ::KAN*, *GFP-MAD3 (HIS3)* | S1G |
| 5143 | BY4742 | *MAT α, mad3Δ::KAN*, *GFP-MAD3 (HIS3)* | S1G |
| 5453 | YEF473 | *MAT a*, *SPC25*-mCherry-*HYG*, *pYGFP-MAD3* | 1E |
| 5454 | YEF473 | *MAT α*, *SPC25*-mCherry-*HYG*, *pYGFP-MAD3* | 1E |
| 4989 | BY4741 | *MAT a, fpr1Δ, MPS1-FRB:KAN, BUB3-2xFKBP12-HYG* | S1B |
| 6178 | YEF473 | *bub1Δ::TRP1::bub1^-abba^ (F490A,V492A,F493A,N495A) (KAN), Mad1-mCherry-HYG, nup60Δ::TRP1, Ndc80-GFP-NAT* | S1C |
| 6179 | YEF473 | *bub1Δ::TRP1::bub1^-abba^ (F490A,V492A,F493A,N495A) (KAN), Mad1-mCherry-HYG, nup60Δ::TRP1, Ndc80-GFP-NAT* | S1C |
| 3781 | YEF473 | *MAT a, spc105Δ::NAT, SPC105*^222::GFP^ *(LEU2), nup60Δ::Trp1, MAD1-mcherry-HYG* | S1C, |
| 5713 | YEF473 | *MAT a, bub1Δ::TRP1::*bub1^T455A,T485A^-*2XFKBP12 (KAN),MAD1*-mCherry-*HYG, nup60Δ::TRP1, SPC25-GFP-HIS3* | S1C |
| 5716 | YEF473 | *MAT* a, *bub1Δ::TRP1::*bub1^T485A,T509A,T518A^-*2XFKBP12 (KAN),MAD1*-mCherry-*HYG*, *nup60Δ::TRP1*, *NNF1-GFP-NAT* | S1C |
| 5717 | YEF473 | *MAT* a, *bub1Δ::TRP1::*bub1^T485A,T509A,T518A^-*2XFKBP12 (KAN),MAD1*-mCherry-*HYG*, *nup60Δ::TRP1*, *NNF1-GFP-NAT* | S1C |
| 5337 | YEF473 | *MAT A, fpr1Δ, bub1Δ::TRP1* | S1D, S2D |
| 6166 | BY4743 | *fpr1Δ, bub1Δ::NAT::bub1-15A -ymCherry (KAN), MAD1-2xFKBP12-HIS3, NDC80-GFP-NAT* | S1E |
| 6167 | BY4743 | *fpr1Δ, bub1Δ::NAT::bub1-15A -ymCherry (KAN), MAD1-2xFKBP12-HIS3, NDC80-GFP-NAT* | S1E |
| 6143 | YEF473 | *MAT a, fpr1Δ, bub1Δ::TRP1::*bub1-15A*-ymCherry (KAN), Mad1-2XFKBP12-HYG, Mps1-FRB-NAT* | S1F |
| 6144 | YEF473 | *MAT a, fpr1Δ, bub1Δ::TRP1::*bub1-15A*-ymCherry (KAN), Mad1-2XFKBP12-HYG, Mps1-FRB-NAT* | S1F |
| 4371 | YEF473 | *MAT a, spc105Δ::NAT*, Spc105^222::GFP^ (*LEU2*), *IPL1*-mCherry-*HYG* | 2A |
| 4372 | YEF473 | *MAT a, spc105Δ::NAT,* Spc105^222::GFP^ (*LEU2*), *IPL1*-mCherry-*HYG* | 2A |
| 5305 | YEF473 | *MAT a, fpr1Δ,* *IPL1-2xFKBP12-HYG*, *GFP*-6XMELT-*FRB (HIS3)* | 2B |
| 5304 | YEF473 | *MAT a, fpr1Δ, IPL1-2xFKBP12-HYG*, *GFP-*6XMELT- *FRB (HIS3)* | 2B |
| 5205 | YEF473 | *MAT a, fpr1Δ*, *IPL1-FRB-KAN*, *BUB1-2xFKBP12-HIS3* | 2B |
| 5206 | YEF473 | *MAT a, fpr1Δ, IPL1-FRB-KAN, BUB1-2xFKBP12-HIS3* | 2B |
| 5224 | YEF473 | *fpr1Δ, IPL1-FRB-KAN, BUB1-2xFKBP12-HIS3, mad1Δ::NAT* | 2B |
| 5281 | YEF473 | *MAT a, fpr1Δ, IPL1-FRB-KAN, MAD1-2xFKBP12-HYG* | 2B |
| 5282 | YEF473 | *MAT a, fpr1Δ, IPL1-FRB-KAN, MAD1-2xFKBP12-HYG* | 2B |
| 6020 | YEF473 | *MAT a, fpr1Δ, IPL1-FRB-KAN, BUB1-2xFKBP12-HIS3*, Spc105^455::GFP^ (*LEU2*) | 2B |
| 6021 | YEF473 | *MAT a, fpr1Δ, Ipl1-FRB-KAN*, *BUB1*-2x*FKBP12*-*HIS3*, Spc105^455::GFP^ (*LEU2*) | 2B |
| 6022 | YEF473 | *MAT a, fpr1Δ, IPL1-FRB-KAN, MAD1-2xFKBP12-HYG*, Spc105^455::GFP^ (*LEU2*) | 2B |
| 6023 | YEF473 | *MAT a, fpr1Δ, IPL1-FRB-KAN, MAD1-2xFKBP12-HYG,* Spc105^455::GFP^ (*LEU2*) | 2B |
| 6230 | YEF473 | *MAT α, fpr1Δ, BUB1-1xFKBP12-HIS3, DSN1-ymCherry-HYG, IPL1-FRB-GFP (URA3)* | 2C |
| 6231 | YEF473 | *MAT α, fpr1Δ, BUB1-1xFKBP12-HIS3, DSN1-ymCherry-HYG, IPL1-FRB-GFP (URA3)* | 2C |
| 6232 | YEF473 | *MAT α, fpr1Δ, BUB1-1xFKBP12-HIS3, DSN1-ymCherry-HYG, ipl1-^K133R^-FRB-GFP (URA3)* | 2C |
| 6233 | YEF473 | *MAT α, fpr1Δ, BUB1-1xFKBP12-HIS3, DSN1-ymCherry-HYG, ipl1-^K133R^-FRB-GFP (URA3)* | 2C |
| 6260 | YEF473 | *MAT a, fpr1Δ, Ipl1-FRB-NAT, GFP-bub1^368-609^-2xFKBP12 (HIS3), MAD1-5xFLAG-KAN, NDC80-ymCherry-HYG* | 2C, S2A |
| 6261 | YEF473 | *MAT a, fpr1Δ, Ipl1-FRB-NAT, GFP-bub1^368-609^-2xFKBP12 (HIS3), MAD1-5xFLAG-KAN, NDC80-ymCherry-HYG* | 2C, S2A |
| 5313 | YEF473 | *MAT a, fpr1Δ, Ipl1-FRB:NAT, bub1Δ::TRP1::*bub1-abba^(F490A,V492A,F493A,N495A)^-*2XFKBP12(KAN)* | 2D |
| 6071 | BY4743 | *MAT a, fpr1Δ, bub1::NAT::*bub1^T453A,T455A^-*2XFKBP12 (KAN), IPL1-FRB-NAT* | 2D |
| 6072 | BY4743 | *MAT a, fpr1Δ, bub1::NAT::*bub1^T453A,T455A^-*2XFKBP12 (KAN), IPL1-FRB-NAT* | 2D |
| 6074 | BY4743 | *MAT a, fpr1Δ, bub1::NAT::*bub1^T455A,T485A^-*2XFKBP12 (KAN), IPL1-FRB-NAT* | 2D |
| 6075 | BY4743 | *MAT a, fpr1Δ, bub1::NAT::*bub1^T455A,T485A^-*2XFKBP12 (KAN),* *IPL1-FRB-NAT* | 2D |
| 6256 | YEF473 | *MAT a, fpr1Δ, Ipl1-FRB-NAT, GFP-bub1-abba^368-608, F490A,V492A,F493A,N495A^-2xFKBP12 (HIS3), MAD1-5xFLAG-KAN* | 2D |
| 6257 | YEF473 | *MAT a, fpr1Δ, Ipl1-FRB::NAT, GFP-bub1-abba^368-608, F490A,V492A,F493A,N495A^-2xFKBP12 (HIS3), MAD1-5xFLAG-KAN* | 2D |
| 5958 | YEF473 | *fpr1Δ, MAD1-2XFKBP12-HIS3, IPL1-FRB-NAT, bub1Δ::TRP1* | 2E |
| 5929 | YEF473 | *MAT a*, *fpr1Δ, IPL1*-*FRB-NAT*, *bub1Δ::TRP1::*bub1-ABBA^(F490A,V492A,F493A,N495A)^ (*KAN*), *MAD1*-2x*FKBP12*-*HIS3* | 2E |
| 5583 | BY4741 | *MAT a, fpr1Δ, IPL1-FRB-KAN, MAD1-2xFKBP12-HYG,* bub1^T485A, T509A, T518A^ *-9xMYC (HIS3)* | 2E |
| 5116 | YEF473 | *MAT a/α, SPC105/spc105Δ::NAT,* spc105^RASA (V76, F78::A), 222::GFP^ (*URA3*), *BUB1/bub1Δ::TRP1::*bub1-abba^(F490A,V492A,F493A,N495A)^ (*KAN*) | 2F |
| 5117 | YEF473 | *MAT a/α*, *SPC105/spc105Δ::NAT*, *spc105*^RASA (V76, F78::A), 222::GFP^ (*URA3*), *BUB1/bub1Δ::TRP1::*bub1-abba^(F490A,V492A,F493A,N495A)^ (*KAN*) | 2F |
| 5132 | YEF473 | *MAT a, spc105Δ::NAT,* *spc105*^RASA (V76, F78::A), 222::GFP^ (*URA3*), *bub1Δ::TRP1::*bub1-abba^(F490A,V492A,F493A,N495A)^ (*KAN*) | 2F |
| 5133 | YEF473 | *MAT α, spc105Δ::NAT*, *spc105*^RASA (V76, F78::A), 222::GFP^ (*URA3*), *bub1Δ::TRP1*::bub1-abba^(F490A,V492A,F493A,N495A)^ (*KAN*) | 2F |
| 5140 | YEF473 | *MAT a, bub1Δ::TRP1::*bub1-abba^(F490A,V492A,F493A,N495A)^ (*KAN*) | 2F |
| 5141 | YEF473 | *MAT α, bub1Δ::TRP1::*bub1-abba^(F490A,V492A,F493A,N495A)^ (*KAN*) | 2F |
| 6245 | YEF473 | *MAT a, fpr1Δ, TetR-GFP (LEU2), CENIV-TetO-URA3, IPL1-FRB-NAT, SPC98-mCherry-HIS3, BUB1-2xFKBP12-HYG* | S2B |
| 6246 | YEF473 | *MAT a, fpr1Δ, TetR-GFP (LEU2), CENIV-TetO:Ura, IPL1-FRB-NAT, SPC98-mCherry-HIS3, BUB1-2xFKBP12-HYG* | S2B |
| 5436 | YEF473 | *MAT a, fpr1Δ, Ipl1-FRB:NAT, bub1Δ*::*TRP1*::bub1^T453A^-*2XFKBP12 (KAN)* | S2C, S2D, S2G |
| 5437 | YEF473 | *MAT a, fpr1Δ, Ipl1-FRB:NAT, bub1Δ::TRP1*::bub1^T453A^-*2XFKBP12 (KAN)* | S2C, S2D, S2G |
| 5622 | YEF473 | *MAT a, fpr1Δ, Ipl1-FRB:NAT, bub1Δ::TRP1::*bub1^T509A, T518A^-*2XFKBP12 (KAN)* | S2C, S2G |
| 5623 | YEF473 | *MAT a, fpr1Δ, Ipl1-FRB:NAT, bub1Δ::TRP1::*bub1^T509A, T518A^-*2XFKBP12 (KAN)* | S2C, S2G |
| 6086 | BY4743 | *MAT a, fpr1Δ, bub1::NAT::*bub1^T485A,T509A,T518A^-*2XFKBP12 (KAN), Ipl1-FRB-NAT* | S2C, S2G |
| 6087 | BY4743 | *MAT a, fpr1Δ, bub1::NAT::*bub1^T485A,T509A,T518A^-*2XFKBP12 (KAN), Ipl1-FRB-NAT* | S2C, S2G |
| 6063 | BY4743 | *MAT a,* *fpr1Δ, bub1::NAT::*bub1^T453A,T455A^-*2XFKBP12 (KAN)* | S2D |
| 6064 | BY4743 | *MAT a,* *fpr1Δ, bub1::NAT::*bub1^T453A,T455A^-*2XFKBP12 (KAN)* | S2D |
| 6065 | BY4743 | *MAT* a, *fpr1Δ, bub1::NAT::* bub1*^T455A,T485A^-2XFKBP12 (KAN)* | S2D |
| 6066 | BY4743 | *MAT* a, *fpr1Δ, bub1::NAT::* bub1*^T455A,T485A^-2XFKBP12 (KAN)* | S2D |
| 6082 | BY4743 | *MAT* α, *fpr1Δ, bub1::NAT::*bub1^T485A,T509A,T518A^-*2XFKBP12 (KAN)* | S2D |
| 6083 | BY4743 | *MAT* a, *fpr1Δ, bub1::NAT::*bub1^T485A,T509A,T518A^-*2XFKBP12 (KAN)* | S2D |
| 6088 | BY4743 | *MAT* a, *fpr1Δ, bub1::NAT::*bub1^T509A,T518A^-*2XFKBP12 (KAN)* | S2D |
| 6089 | BY4743 | *MAT* a, *fpr1Δ, bub1::NAT::*bub1^T509A,T518A^-*2XFKBP12 (KAN)* | S2D |
| 6094 | BY4743 | *MAT a,* *fpr1Δ* | S1D, S2D |
| 6145 | YEF473 | *MAT α,* *fpr1Δ, IPL1-FRB-GFP-NAT, BUB1-1xFKBP12-HIS3, mps1Δ::KAN, mps1-as1 (mps1-M516G, CEN, TRP1), DSN1-ymCherry-HYG* | S2E |
| 6146 | YEF473 | *MAT α, fpr1Δ, IPL1-FRB-GFP-NAT, BUB1-1xFKBP12-HIS3, mps1Δ::KAN, mps1-as1 (mps1-M516G, CEN, TRP1), DSN1-ymCherry-HYG* | S2E |
| 5624 | YEF473 | *MAT a/MAT α, SPC105/spc105Δ::NAT*, spc105^RASA (V76, F78::A) 222::GFP^ (*URA3*), *BUB1/bub1Δ::TRP1::*bub1^T485A,T509A,T518A^-*2XFKBP12 (KAN)* | S2F |
| 5625 | YEF473 | *MAT a/MAT α, SPC105/spc105Δ::NAT*, spc105^RASA (V76, F78::A) 222::GFP^ (*URA3*), *BUB1/bub1Δ::TRP1::*bub1^T455A, T485A^ -*2XFKBP12 (KAN)* | S2F |
| 6069 | BY4743 | *MAT a, fpr1Δ, bub1::NAT::*bub1^T453A,T455A^-*2XFKBP12 (KAN),* *MPS1-FRB-NAT* | S2G |
| 6070 | BY4743 | *MAT a, fpr1Δ, bub1::NAT::*bub1^T453A,T455A^-*2XFKBP12 (KAN), MPS1-FRB-NAT* | S2G |
| 6073 | BY4743 | *MAT a, fpr1Δ, bub1::NAT::*bub1^T455A,T485A^-*2XFKBP12 (KAN), MPS1-FRB-NAT* | S2G |
| 6077 | BY4743 | *MAT a, fpr1Δ, bub1::NAT::*bub1^T455A,T485A^-*2XFKBP12 (KAN), MPS1-FRB-NAT* | S2G |
| 6084 | BY4743 | *MAT a, fpr1Δ, bub1::NAT::*bub1^T485A, T509A, T518A^-*2XFKBP12 (KAN), MPS1-FRB-NAT* | S2G |
| 6085 | BY4743 | *MAT a, fpr1Δ, bub1::NAT::*bub1^T485A, T509A, T518A^-*2XFKBP12 (KAN), MPS1-FRB-NAT* | S2G |
| 5724 | YEF473 | *MAT a, fpr1Δ, Ipl1-FRB-NAT, bub1Δ::TRP1::bub1-15A-2XFKBP12 (KAN)* | S2G |
| 5725 | YEF473 | *MAT a, fpr1Δ, Ipl1-FRB-NAT, bub1Δ::TRP1::bub1-15A-2XFKBP12 (KAN)* | S2G |
| 5473 | YEF473 | *MAT a, fpr1Δ, Mps1-FRB-NAT, bub1Δ::TRP1::bub1^T453A^-2XFKBP12 (KAN)* | S2G |
| 5474 | YEF473 | *MAT a, fpr1Δ, Mps1-FRB-NAT, bub1Δ::TRP1::bub1^T453A^-2XFKBP12 (KAN)* | S2G |
| Yeast strains discussed in the text but not in the figures | | | |
| 3802 |  | *MAT a, -leu2, -trp1,-ura3, -his3, -ade2* | Not applicable |
| 6247 | YEF473 | *MAT a, fpr1Δ, IPL1-FRB-NAT, GFP-*bub1^368-609^*-2xFKBP12 (HIS3), MET3pr-3xHA-CDC20 (URA3)* |  |
| 6249 | YEF473 | *MAT a, fpr1Δ, IPL1-FRB-NAT, GFP-*bub1^368-609, F490A,V492A, F493A, N495A^*-2xFKBP12 (HIS3), MET3pr-3xHA-CDC20 (URA3)* |  |
| 6250 | YEF473 | *MAT a, fpr1Δ, IPL1-FRB-NAT, GFP-*bub1^368-609, F490A, V492A, F493A, N495A^ *-2xFKBP12 (HIS3), MET3pr-3xHA-CDC20 (URA3)* |  |
| 5259 | YEF473 | *MAT a/MAT α, SPC105/spc105Δ::NAT*, spc105^RASA (V76, F78::A), 222::GFP^ (*URA3*), *CDC20/cdc20Δ::TRP1::*CDC20^S24A^(*KAN*) |  |
| 5261 | YEF473 | *MAT a/MAT α, SPC105/spc105Δ::NAT*, spc105^RASA (V76, F78::A), 222::GFP^ (*URA3*), *CDC20/cdc20Δ::TRP1::*CDC20^S24A,^ ^S46A, S52A, S62A, S88A, S89A^ (*KAN*) |  |
| 5350 | YEF473 | *MAT a/MAT α, SPC105/spc105Δ::NAT*, spc105^RASA (V76, F78::A), 222::GFP^ (*URA3*), *CDC20/cdc20Δ::TRP1::*CDC20^S602A^ (*KAN*) |  |
| 5351 | YEF473 | *MAT a/Mat α, SPC105/spc105Δ::NAT*, spc105^RASA (V76, F78::A), 222::GFP^ (*URA3*), *CDC20/cdc20Δ::TRP1::*CDC20^602-605::AAAA^ (*KAN*) |  |
| 5355 | YEF473 | *MAT a/Mat α, SPC105/spc105Δ::NAT*, spc105^RASA (V76, F78::A), 222::GFP^ (*URA3*), *CDC20/cdc20Δ::TRP1::*CDC20^S24A,^ ^S46A, S52A, S62A, S88A, S89A, 602-605::AAAA^ (*KAN*) |  |
| 5415 | YEF473 | *MAT a/Mat α, SPC105/spc105Δ::NAT*, spc105^RASA (V76, F78::A), 222::GFP^ (*URA3*), *CDC20/cdc20Δ::TRP1::*CDC20^S229A^ (*KAN*) |  |

| Plasmid (pAJ#) | Origin | Parent | Description |
| --- | --- | --- | --- |
| pAJ147 | Biggins lab | pRS314 | *mps1-as1 (*mps1-M516G*), (TRP1, CEN)* |
| pAJ332 | Joglekar lab | pRS305 | *SPC105*pr+Spc105-6A^(T149A, T172A, T211A, T235A, T284A, T313A)^-12X*MYC*+*trSPC105 (LEU2)* |
| pAJ351 | Joglekar lab | pRS305 | *HIS3*pr*-GFP(S65T)-* Spc105^120-329^-FRB- SV40 NLS *(LEU2)* |
| pAJ387 | Joglekar lab | pRS305 | *HIS3*pr*-GFP(S65T)-* Spc105^120-128^-FRB- SV40 NLS *(LEU2)* |
| pAJ419 | This study | pRS305 | *SPC105*pr+*Spc105^455::GFP^+ trSPC105 (LEU2)* |
| pAJ551 | Joglekar lab | pRS305 | *HIS3*pr*-GFP(S65T)-*Spc105^2-329^-FRB-SV40 NLS *(LEU2)* |
| pAJ591 | Biggins lab | pRS303 | bub1^T485A, T509A, T518A^ *-*9xMYC *(HIS3)* |
| pAJ808 | This study | pSK954 | *BUB1*pr+bub1^-abba (F490A, V492A, F493A, N495A)^ +*BUB1*Tr (*KAN*) |
| pAJ821 | This study | pSK954 | *BUB1*pr+bub1^-abba (F490A,V492A,F493A,N495A)^+yeast mCherry +*BUB1*Tr (*KAN*) |
| pAJ827 | This study | pRS303 | GFP*-Mad1(CTD)-2xFKBP12 (HIS3)* |
| pAJ831 | This study | pSK954 | *BUB1*pr+bub1^-abba (F490A,V492A,F493A,N495A)^+2X*FKBP12* +*BUB1*Tr (*KAN*) |
| pAJ847 | This study | pSK954 | *BUB1*pr+bub1^T453A^+2X*FKBP12* +*BUB1*Tr (*KAN*) |
| pAJ852 | Joglekar lab | pSK954 | *BUB1*pr+bub1^T453A, T455A^+2X*FKBP12* +*BUB1*Tr (*KAN*) |
| pAJ857 | This study | pRS303 | *GFP-Mad1(CTD)-4A^(T624A+T704A+T737A+T739A)^-2xFKBP12 (HIS3)* |
| pAJ873 | This study | pSK954 | *BUB1*pr+bub1^T485A, T509A, T518A^+2X*FKBP12* +*BUB1*Tr (*KAN*) |
| pAJ874 | This study | pSK954 | *BUB1*pr+bub1^T455A, T485A^+2X*FKBP12* +*BUB1*Tr (*KAN*) |
| pAJ875 | This study | pSK954 | *BUB1*pr+bub1^T509A, T518A^+2X*FKBP12* +*BUB1*Tr (*KAN*) |
| pAJ886 | This study | pSK954 | *BUB1*pr+bub1^T485A, T486A, T487A, T488A, T509A,T518A, S537A,S539A,S540A, T541A, T555A,T556A,T558A,T566A, S578A^+2X*FKBP12* +*BUB1*Tr (*KAN*) |
| pAJ896 | Joglekar lab | pSK954 | *BUB1*pr+bub1^T453A, 455A^+2X*FKBP12* +*BUB1*Tr (*KAN*) |
| pAJ923 | This study | pSK954 | *BUB1*pr+bub1^T485A, T486A, T487A, T488A, T509A,T518A, S537A,S539A,S540A, T541A, T555A,T556A,T558A,T566A, S578A^+yeast mCherry+*BUB1*Tr (*KAN*) |
| pAJ932 | This study | pAFS144 | *HIS3pr+GFP-*bub1^368-609^*-2xFKBP12 (HIS3)* |
| pAJ934 | This study | pAFS144 | *HIS3pr+GFP-*bub1^368-609, F490A,V492A,F493A,N495A^*-2xFKBP12 (HIS3)* |
| pAJ940 | This study | pRS306 | *IPL1pr+*ipl1^K133R^*-FRB-GFP+IPL1Tr (URA3)* |
| pAJ941 | This study | pRS306 | *IPL1pr+IPL1-FRB-GFP+IPL1Tr (URA3)* |
| **Plasmids discussed in the text for yeast study but not in the figures** | | | |
| pAJ810 | This study | pSK954 | *CDC20*pr*+CDC20^S24A^+CDC20*Tr *(KAN)* |
| pAJ816 | This study | pSK954 | *CDC20*pr*+CDC20^S24A, S46A, S52A, S62A, S88A, S89A^+CDC20*Tr *(KAN)* |
| pAJ837 | This study | pSK954 | *CDC20*pr*+CDC20^S24A, S46A, S52A, S62A,S88A,S89A, 602-605::AAAA^+CDC20*Tr *(KAN)* |
| pAJ838 | This study | pSK954 | *CDC20*pr*+CDC20^602A^+CDC20*Tr *(KAN)* |
| pAJ839 | This study | pSK954 | *CDC20*pr*+CDC20 ^602-605::AAAA^+CDC20*Tr *(KAN)* |
| pAJ850 | This study | pSK954 | *CDC20*pr*+CDC20^S229A^+CDC20*Tr *(KAN)* |
| pAJ935 | This study | pGBD_C1 | pGBD_C1+bub1^368-609^ |
| pAJ936 | This study | pGBD_C1 | pGBD_C1+bub1^368-609, F490A, V492A, F493A, N495A^ |
| pAJ937 | This study | pGAD_C1 | pGAD_C1*+CDC20 (LEU2)* |
| pAJ950 | This study | pGBD_C1 | pGBD_C1+bub1^451-520^ |
| pAJ951 | This study | pGBD_C1 | pGBD_C1+bub1^451-520, F490A, V492A, F493A, N495A^ |
| pAJ952 | This study | pGBD_C1 | pGBD_C1+ bub1^N-609^ |
| pAJ953 | This study | pGBD_C1 | pGBD_C1+ bub1^N-609, F490A, V492A, F493A, N495A^ |
| **Plasmids used for constructing stable HeLa cell lines** | | | |
| pERB131 | Lampson lab |  | Mis12-GFP-FKBP12x3/Inducible mcherry-Mps1 |
| pPS88 | This study | pERB131 | Bub1^271-620^-mNeonGreen-2xFkbp12/Frb-mCherry-Aurora B^61-344^ (kinase domain) |
| pPS95 | This study | pERB131 | mNeonGreen-2xFkbp12/Frb-mch-INCENP^818-918^ |
| pPS94 | This study | pERB131 | M3-M3-mNeonGreen-2xFkbp12/Frb-mcherry-INCENP^818-918^ |
| pPS96 | This study | pERB131 | Mad1L1^479-725^-mNeonGreen-2xFkbp12/Frb-mCherry-INCENP^818-918^ |
| pPS108 | This study | pERB131 | Bub1^225-620^-mNeonGreen-2xFkbp12/Frb-mCherry-INCENP^818-918^ |
| pPS69 | This study | pERB131 | Bub1^271-620^-mNeonGreen-2xFkbp12/Frb-mCherry-Mps1 kinase domain |
| pPS99 | This study | pERB131 | Bub1^271-620^-mNeonGreen-2xFkbp12/Frb-mCherry-INCENP^818-918^ |
| pPS85 | This study | pERB131 | Bub1^271-523^-mNeonGreen-2xFkbp12/Frb-mCherry-Mps1 kinase domain |
| pPS98 | This study | pERB131 | Bub1^271-523^-mNeonGreen-2xFkbp12+Frb-mCh-INCENP^818-918^ |
| pMF1502 | Arshad Desai lab |  | AAVS1 homology arms, Histone H2B-mRFP1.3(K55E) |
| pX330 | Addgene |  | AAVSI T2 CRISPR gRNA, SpCas9-mNeonGreen |

**Table S3.** Lists of *S. cerevisiae* strains and Plasmids/Recombinant DNA used in this study.
